# Healthcare experience affects pain-specific responses to others’ suffering in the anterior insula

**DOI:** 10.1101/2021.07.01.450687

**Authors:** Corrado Corradi-Dell’Acqua, Christoph Hofstetter, Gil Sharvit, Olivier Hugli, Patrik Vuilleumier

## Abstract

Medical students and professional healthcare providers often underestimate patients’ pain, together with decreased neural responses to pain information in the anterior insula (AI), a brain region implicated in self-pain processing and negative affect. However, the functional significance and specificity of these neural changes remains debated. Across two experiments, we recruited university medical students and emergency nurses to test the role of healthcare experience on the brain reactivity to other’s pain, emotions, and beliefs, using both pictorial and verbal cues. Brain responses to self-pain was also assessed and compared with those to observed pain. Our results confirmed that healthcare experience decreased the activity in AI in response to others’ suffering. This effect was independent from stimulus modality (pictures or texts), but specific for pain, as it did not generalize to inferences about other mental or affective states. Furthermore, representational similarity and multivariate pattern analysis revealed that healthcare experience impacted specifically a component of the neural representation of others’ pain that is shared with that of first-hand nociception, and related more to AI than to other pain-responsive regions. Taken together, our study suggests a decreased propensity to appraise others’ suffering as one’s own, associated with a reduced recruitment of pain-specific information in AI. These findings provide new insights into neural mechanisms leading to pain underestimation by caregivers in clinical settings.

## Introduction

Unrelieved pain is a major problem worldwide, resulting in human suffering and economic costs. In medical practice, pain is difficult to quantify objectively, and it is often assessed indirectly through clinical examination and patients’ self-reports. It is therefore not surprising that healthcare providers may underestimate (Ruben et al., 2015, 2018) (and undertreat, Rupp & Delaney, 2004) patients’ pain, a phenomenon that emerges during university education (Dirupo et al., 2021; Xie et al., 2018), and becomes more pronounced with longer experience in the field (Choinière et al., 1990; Davoudi et al., 2008).

Neuroscience research has investigated extensively the cerebral mechanisms that underlie the appraisal of other’s pain, and has begun to unveil how they are modified by healthcare training. Imaging studies have implicated a widespread brain network, centered around the anterior insula (AI) and dorsal-anterior cingulate cortex (dACC), in the processing and empathizing with other people’s pain (as conveyed by faces, pictures, text, etc.) (Ding et al., 2019; Y. Fan et al., 2011; Jauniaux et al., 2019; Kogler et al., 2020; Lamm et al., 2011; Schurz et al., 2021; Timmers et al., 2018). Remarkably, these activity patterns are highly similar to those measured when pain is experienced directly by oneself (Berluti et al., 2020; Braboszcz et al., 2017; Corradi-Dell’Acqua et al., 2011, 2016; O’Connell et al., 2019; Qiao-Tasserit et al., 2018; Wagner et al., 2020; Zhou et al., 2020), and they are attenuated by those same analgesic procedures that regulate first-hand nociception, such as placebo or hypnosis (Braboszcz et al., 2017; Rütgen et al., 2015, 2021). These results suggest that others’ pain is at least partly processed in an embodied (or empathetic) fashion, that is, by simulating its somatic and affective properties on one’s own body (Bastiaansen et al., 2009; Bernhardt & Singer, 2012).

Critically, medical practitioners and students exhibit lower activity in these regions to the sight of injuries and painful expressions (Chen et al., 2022; Cheng et al., 2007, 2017; Dirupo et al., 2021; Jackson et al., 2017). It has been hypothesized that continuous interaction with severe conditions and injuries might make healthcare providers progressively desensitized towards the sight of pain, possibly due to regulatory processes protecting them from the psychological costs of repeated exposure to suffering (Chen et al., 2022; Gleichgerrcht & Decety, 2014; Vaes & Muratore, 2013).

Recently, however, scholars have underscored that the neural response in the insula and cingulate cortex is not specific for pain, but responds also to a wide range of painless conditions, including various emotional events (Corradi-Dell’Acqua et al., 2011, 2016) and non-affective but intense visual/auditory stimulations (Liang et al., 2019). This lack of specificity challenged embodied interpretations of social cognition and empathy, as it is difficult to disentangle components of neural activity that underlie specific affective states from those coding for supra-ordinal dimensions, such as the unpleasantness, intensity, or salience of an event (Corradi-Dell’Acqua et al., 2016; Rütgen et al., 2021; Sharvit et al., 2020). This problem also concerns studies investigating healthcare training, which could focus on the neural response of pain-specific processes shared between oneself and others (*seeing injured patients hurts me*), or broader mechanisms that signal any emotionally or attentionally salient stimulus (*seeing injured patients captures my attention*).

Here, we reanalysed the data from two independent experiments in which university students at different years of medical school (plus controls from other faculties; cohort 1; Corradi-Dell’Acqua et al., 2011, 2014) and professional healthcare providers (cohort 2, Corradi-Dell’Acqua et al., 2019) underwent highly similar experimental protocol where they were exposed to: (1) nociceptive thermal stimulations to their own hand, and (2) pictures and narratives describing others in pain, as well as in control painless states. This allowed us to assess whether scholarly (cohort 1) and professional (cohort 2) healthcare experience affects the neural responses to others’ pain (Chen et al., 2022; Cheng et al., 2007, 2017; Dirupo et al., 2021; Jackson et al., 2017) and, most critically, whether such influence operates on state-specific representations shared with first-hand nociception or generalizes to other emotional events.

## Methods

### Participants

The present study was carried out on two independent cohorts. Cohort 1 included 43 female students from the University of Geneva. Part of these participants were recruited in previous studies (Corradi-Dell’Acqua et al., 2011, 2014), where they were tested as a unique group, without taking into account the faculty in which they were enrolled. Here, we included only those individuals who fell into the following three groups: students enrolled in the first year of medical school (*Med1*: N = 15, age 18-22 years, mean = 19.8); students enrolled in the fourth year of medical school (*Med4*: N = 14, age 22-29 years, mean = 24.14); and students attending other university faculties or high schools except medicine, infirmary, dentistry, or kinesi-physiotherapy (*Controls*: N = 14, age 19-31 years, mean = 23.42).

Cohort 2 included 30 *Nurses* from the Emergency Department of the University Hospital of Lausanne (females = 17, age range = 24-61, mean = 36.33; post-graduate experience = 4-33 years, mean = 11.03), which were part of the sample in a previous study (Corradi-Dell’Acqua et al., 2019), although their response was never tested as function of their professional (post-graduate) experience.

None of our participants had any history of neurological or psychiatric illness. Written informed consent was obtained from all subjects. The studies were approved by the ethical committee of the University Hospital of Geneva (cohort 1) and by the Ethical Cantonal Commission of Canton Vaud (cohort 2), and conducted according to the declaration of Helsinki.

### Experimental Protocol

Participants from both cohorts underwent three separate experimental sessions in the Magnetic Resonance Imaging (MRI) scanner, followed by a post-scanning rating session. These paradigms involved exposure to others’ pain through pictures of injured hands (*Handedness task*) or through brief narratives (*Cognitive and Affective Theory of Mind task*), as well as the delivery of pain to one’s own body through painful thermal stimulation (*Pain Localizer*). Both the “Handedness” and “Cognitive and Affective Theory of Mind” tasks were extensively described in previous studies (Corradi-Dell’Acqua et al., 2011, 2014) and used in follow-up investigations (Corradi-Dell’Acqua et al., 2019, 2020; Qiao-Tasserit et al., 2018).

#### Handedness Task

Participants from Cohort 1 underwent the same protocol as described in Corradi-Dell’Acqua et al. (2011), whereas participants from Cohort 2 underwent a shorter version of the task (120 instead of 180 trials) similar to Corradi-Dell’Acqua et al. (2019). In both cases, participants saw a randomized sequence of colour pictures (768×768 pixels, corresponding to 14.25°x14.25° degrees of visual angle) of human hands, organized in four categories. The “Painful” category (*PF*) was composed of 60 (cohort 1) or 30 (cohort 2) images depicting hands in painful situations, inferable by either the presence of wounds/burns on the skin and/or an external object (scalpel, syringe, etc.) acting on the skin surface. The negative “Painless” category (*PL*) was composed of 30 pictures of hands in emotionally aversive, but painless situations (hands holding knifes/guns, hands with handcuffs, etc.). For both *PF* and *NPL* stimuli, we also selected neutral control stimuli (*cPF* and *cPL*) that were matched with the previous two categories for hand laterality (right/left) orientation (angular distance from the viewer’s own hand position at rest), and for visual features (presence of objects, human bodies, etc.), but purged from any emotionally-salient (painful, arousing) features. This yielded a 2 x 2 design with STIMULI (*Painful*, *Painless*) and EMOTIONAL AROUSAL (*Negative*, *Neutral*) as factors. All images were equated in luminance and are available under the Open Science Framework at the following link: https://osf.io/8bjmq/.

For each experimental trial, one of the hand stimuli was presented for 2500 ms, followed by an inter-trial interval that ranged from 2500 to 4100 ms (mean and median = 3300 ms) with incremental steps of 320 ms. Participants had to perform a handedness task, i.e., to report if the stimulus depicted a right or left hand by pressing a corresponding key. This task is known to be accomplished by mentally imagining to move one’s own hand until it is aligned with the viewed hand (Corradi-Dell’Acqua et al., 2009), hence favoring an embodied perspective, but it did not make any explicit demand to process the painful or emotional cues in pictures. Participants were instructed to respond as fast as possible and to ignore other image features (e.g., blades, wounds) that were irrelevant to the task. The 4 experimental conditions of this task were presented in a randomized order together with 30 null-events, in which an empty screen replaced the stimuli. All trials were presented in a unique scanning session which lasted about 21 (cohort 1) or 15 (cohort 2) minutes.

#### Cognitive and Affective Theory of Mind Task

Participants from Cohort 1 underwent the same protocol as described in Corradi-Dell’Acqua et al. (2014), whereas participants from Cohort 2 underwent a shorter version of the task (1 instead of 2 sessions). The original version of the paradigm comprised of 36 short French-written narratives describing a person engaged in various situations, followed by questions probing for the reader’s awareness of the protagonist’s pain, emotions, or beliefs in this situation. As a high-level control condition, we included 12 additional stories with no human protagonist but describing physical entities with changing properties on visual maps or photographs (photos). This database was validated by an independent group of 40 female participants (age: 19–54 years). The full list of narratives, and results from the validation study are reported in Corradi-Dell’Acqua et al. (2014).

The task was organized in experimental sessions containing a random sequence of 24 narratives (6 per conditions) of about 10 minutes. In cohort 1, participants underwent two sessions, covering the full set of 48 scenarios, whereas cohort 2 underwent only one session of 24 scenarios. Within each session, the scenarios were presented for 12 seconds, followed by a judgment epoch of 5 seconds. During the judgment phase, a question was presented together with two possible answers, each located on a different side of the screen. Participants made responses by pressing one of two possible keys, corresponding to the side of the answer they believed to be correct. The position of the correct response on the screen was counterbalanced across narratives. Judgments were followed by an inter-trial interval of 10 seconds.

Importantly, in the “judgment” stage, participants were asked to evaluate only one dimension (beliefs, emotions, pain), but it is likely that during the reading stage, the “scenarios” could elicit spontaneous considerations about multiple dimensions at the time (e.g., a story about someone’s pain often implies thoughts about other emotional reactions and mental states). As a reliable estimate of the scenarios’ likelihood to elicit inferences about each dimension, we used the scores from a previous validation study on the same story database, described in Corradi-Dell’Acqua et al. (2014).

#### Pain Localizer

In this task, participants received either noxious or non-noxious thermal stimulations to their hand palm, delivered by using a computer controlled thermal stimulator with an MRI compatible 25×50 mm fluid cooled Peltier probe (MSA Thermotest, SOMEDIC Sales AB, Sweden). Unlike the other two tasks, the Pain Localizer protocol used different settings in the two cohorts. In cohort 1 thermal events were delivered in two separate runs, involving stimulation of the right or left palm, respectively. Each run comprised ten trials, five with a noxious temperature and five with a non-noxious temperature. Each trial was organized into two consecutive thermal shifts, each lasting 9 seconds (3 seconds of temperature increase, 3 seconds of plateau, and 3 seconds of temperature decrease), followed by an inter-trial interval of 18 seconds. In cohort 2, thermal stimuli were delivered in a single run, with stimulation on the dominant palm. The run comprised twelve trials, six with a noxious temperature and six with a non-noxious temperature. Trials were organized in a unique thermal shift lasting approximately 9 seconds (3 second of temperature increase, 2 seconds of plateau, and 3 seconds of decrease), followed by an inter-trial interval ranging between 9-14.5 sec (average 11.75). In both cohorts, a visual cue (identical for noxious and non-noxious shifts) informed participants of the upcoming shift. The non-noxious temperature was fixed to 36°C (cohort 1) or 38°C (cohort 2). The noxious temperature varied on a participant-by-participant basis and ranged between 41-52°C (cohort 1: average = 46.42°C, *Hot – Warm* difference = 10.89°C; cohort 2: 49.31°C, *Hot – Warm* = 11.31°C). In cohort 1, this temperature was selected through an ascending method of limits (see Corradi-Dell’Acqua et al., 2011; for more details), whereas we used a double random staircase algorithm in cohort 2 (Antico et al., 2018; Corradi-Dell’Acqua et al., 2019; Sharvit et al., 2018, 2020).

#### Post-Scanning rating session

After scanning, participants were asked to rate each of the stimuli from the “Handedness” task in terms of familiarity (“*how much is the content described in this picture familiar to you?*”), emotional intensity (“*how intense is the emotion triggered by this image?*”), emotional valence (*“does this image elicit positive or negative emotions?”*), and pain (“*how intense is the pain felt by the hand depicted on this image?*”). The ratings were divided in four blocks, one for each question, during which all stimuli were rated on a Likert scale ranging from 1 to 10 (with the exception of valence for which a Likert scale from -4 to +4 was used). To avoid habituation biases due to the presentation of the same stimuli four times, the order of the blocks and the order of the stimuli within each block were randomized across participants. See previous reports for more details about the rating session (Corradi-Dell’Acqua et al., 2011).

### Scanning Procedure

Participants lay supine with their head fixated by firm foam pads. Stimuli were presented using E-Prime 2.0 (Psychology Software Tools, Inc.) on a LCD projector (CP-SX1350, Hitachi, Japan) outside the scanner bore, subtending about 14.25° (vertical) x 19° degrees of visual angle. Participants saw the monitor through a mirror mounted on the MR headcoil. Key-presses were recorded on an MRI-compatible bimanual response button box (HH-2×4-C, Current Designs Inc., USA). During the "Pain Localizer", the button box was replaced by the thermode Peltier probe attached on participant’s palm. The order of each task was counterbalanced across participants. In cohort 2, these were intermingled with other paradigms which are described elsewhere (Corradi-Dell’Acqua et al., 2019).

### Data Processing

Our main goal was to test whether behavioral and neural responses associated with our task could be influenced by the degree of healthcare experience. Given that the two cohorts were acquired in non-identical settings, we privileged a separate analysis of each dataset, in the attempt to identify effects of experience that were independently observable and replicable in each group. In cohort 1, the effect of healthcare experience was tested through a between-subject factor GROUP, comprising Controls, Med1, and Med4 students. In cohort 2, instead, we modeled the years of post-graduate EXPERIENCE as a between-subjects covariate of interest.

In a follow-up analysis, limited to parts of the data that were comparable across the two datasets, we combined the two cohorts together to assess the effect of healthcare experience in terms of a four-level GROUP factor (Controls, Med1, Med4, Nurses), with AGE included as a nuisance variable of no interest. In particular, as cohort 1 comprised only women, we included only the 17 female nurses from cohort 2. Furthermore, as cohort 2 underwent a shorter version of the paradigms (see above), we considered only a selection of the data from cohort 1 in order to match the two experiments in terms of power. For the “Handedness” task, we included only those 120 trials (out of 180) which were used in both experiments. For the “Cognitive and Affective Theory of Mind” task, we used only the first run in chronological order. Given the substantial differences associated with the “Pain Localizer”, all measures involving this specific task were not compared across cohorts.

#### Behavioral Data

Data from the "Handedness" task were analyzed through a mixed models schema. Single trial Response Times of correct responses, and post-scanning ratings, were fed in a Linear Mixed Model, whereas for Accuracy values we used a Generalized Linear Mixed Model with binomial distribution and Laplace approximation. We organized the four conditions of interest (*PF*, *PL*, *cPF*, *cPL*) into two orthogonal within-subject factors of STIMULI (*Painful, Painless*) and EMOTIONAL AROUSAL (*Negative, Neutral*). Similarly, in the "Cognitive and Affective Theory of Mind" task, single trial Response Times of correct responses and Accuracy values were fed in a (Generalized) Linear Mixed Model with STORY CATEGORY (*Beliefs*, *Emotion*, *Pain*, *Photos*) as unique within-subject factor. In all models, we tested the effects of healthcare experience, either as a between-subject factor GROUP (e.g., cohort 1), or as a covariate of interest describing the number of years of post-graduate EXPERIENCE (e.g., cohort 2). In all models, participants’ identity was specified as random factor, with random intercept and slope for the within-subject factors (and, where relevant, the interaction thereof). The experimental material (i.e., specific pictures used in the “Handedness Task” or specific scenarios from the "Cognitive and Affective Theory of Mind" task) was modeled as an additional random factor, with random intercept and slope for GROUP or post-graduate EXPERIENCE (cohort 2). For linear mixed models, the significance of the parameter estimates was assessed though the Satterthwaite approximation of the degrees of Freedom. The analysis was run as implemented in the *lmerTest* package (Kuznetsova et al., 2015) of R 4.2.1 software (https://cran.r-project.org/). All data, variables of interest (including information about material items in each subject/trial), and analysis scripts are available under the Open Science Framework (https://osf.io/8bjmq/).

#### Imaging processing

##### Data Acquisition

A Siemens Trio 3-T whole-body scanner was used to acquire both T1- weighted anatomical images and gradient-echo planar T2*-weighted MRI images with blood oxygenation level dependent (BOLD) contrast. For cohort 1, the scanning sequence was a trajectory-based reconstruction sequence with a repetition time (TR) of 2100 msec, an echo time (TE) of 30 msec, a flip angle of 90 degrees, in-plane resolution 64×64 voxels (voxel size 3 x 3 mm), 32 slices, a slice thickness of 3 mm, with no gap between slices. For cohort 2, we used a multiplex sequence (Feinberg et al., 2010), with TR = 650 ms, TE = 30 ms, flip angle = 50°, 36 interleaved slices, 64 x 64 in-slice resolution, 3 x 3 x 3 mm voxel size, and 3.9 mm slice spacing. The multiband accelerator factor was 4, and parallel acquisition techniques (PAT) was not used.

##### Preprocessing

Statistical analysis was performed using the SPM12 software (http://www.fil.ion.ucl.ac.uk/spm/). For each subject, all functional images were fed to a preprocessing pipeline involving realignment (to correct for head movement), unwrapping (to account for geometric distortions related to magnetic field inhomogeneity), slice-time correction (to account for temporal delays within the acquisition of a whole brain volume), and normalization to a template based on 152 brains from the Montreal Neurological Institute (MNI) with a voxel-size resolution of 2 x 2 x 2 mm. Finally, the normalized images were smoothed by convolution with an 8 mm full-width at half-maximum Gaussian kernel.

##### First-Level analysis

Preprocessed images from each task were analyzed using the General Linear Model (GLM) framework implemented in SPM, as in previous studies using the same paradigms (Corradi-Dell’Acqua et al., 2011, 2014, 2019; Qiao-Tasserit et al., 2018). For the “Handedness Task”, trial time onsets from each of the four conditions of interest were modelled with a delta function. For each condition we also included an additional vector in which individual Response Times were modelled parametrically (Corradi-Dell’Acqua et al., 2011, 2019; Qiao-Tasserit et al., 2018). For the “Pain Localizer”, each thermal stimulation was modelled based on the time during which temperature was at plateau.

Finally, for the "Cognitive and Affective Theory of Mind Task" we ran the same model used for Corradi-Dell’Acqua et al. (2014). In particular, we used a boxcar function for the 12 s long blocks during which a scenario was presented, separately from the subsequent 5 s long blocks during which the judgment took place. We modelled four “judgment” vectors, one for each of the four kinds of stories (*Beliefs*, *Emotion*, *Pain*, *Photos*), whereas we modelled only one “scenario” vector, in which all stories were treated as a unique condition. The latter were complemented by four further parametric regressors, one referring to the number of characters in the scenarios, and the other three describing the likelihood to elicit mental attributions beliefs, emotions, and pain, respectively, based on previous validation of the database (Corradi-Dell’Acqua et al., 2014). The scenarios of the photo stories, which had no human protagonists and therefore were not qualified by these dimensions, were associated with an artificial value of 0. To avoid biases related to the order of parametric predictors, and to ensure that each effect was uniquely interpretable, the story epochs were modelled after removing the serial orthogonalization option from SPM default settings. Text-length was also controlled for the “judgment” blocks by including a parametric regressor which, however, made no distinction between the four kinds of stories.

For all tasks, we accounted for putative habituation effects of neural responses in each condition by using the time-modulation option implemented in SPM, which creates a regressor in which the trial order is modulated parametrically. Furthermore, each regressor was convolved with a canonical hemodynamic response function and associated with its first order temporal derivative. To account for movement-related variance, we included six differential movement parameters as covariates of no interest. Low-frequency signal drifts were filtered using a cut-off period of 128 sec. In cohort 1, serial correlation in the neural signal were accounted for through first-order autoregressive model AR(1). In cohort 2, we used instead an exponential covariance structure, as implemented in the ‘FAST’ option of SPM12 for rapid sequences. Global scaling was applied, with each fMRI value rescaled to a percentage value of the average whole-brain signal for that scan.

##### Second-level Analyses

Functional contrasts, comparing differential parameter estimate images associated in one experimental condition *vs.* the other, were then fed in a second level model. These included a one-sample *t*-test testing overall effects across all subjects. For cohort 2, postgraduate EXPERIENCE was included as covariate of interest. For the analysis of cohort 1, and for the comparison between cohort 1 & 2, we also assessed GROUP differences through a one-way ANOVA design. Within these designs, we identified significant effects through Threshold-Free Cluster Enhancement (TFCE) approach, which allows for the identification of a combined voxel-cluster statistics through non-parametric permutation approach (S. M. Smith & Nichols, 2009). This analysis was carried out using the TFCE toolbox for SPM12 (http://dbm.neuro.uni-jena.de/tfce) with 5000 permutations.

##### Volume of interest analysis

In addition to the whole-brain analysis, we also constrained our hypothesis by focusing on voxels from brain regions of theoretical interest. In particular, for contrasts probing individual ability at assessing others’ pain, we defined our volume of interest through the Brainnetome Atlas that provides connectivity-based parcellation of human brain into 246 subregions (L. Fan et al., 2016). In particular, we focused on the “core” pain empathy network which involves bilateral AI and dACC. We therefore created an AI-dACC mask, defined as bilateral cingulate region 3 (corresponding to the pregenual portion of the anterior cingulate cortex) and insular regions 2 and 3 (corresponding approximately to the anterior agranular insular cortex).

##### Vicarious pain signatures

We submitted data from the “Handedness” task to two multivariate neural models predictive of vicarious pain from brain activity. To maximize the comparability with our experiment, we choose models derived from previous datasets in which individuals were shown pictures of limb injuries (Krishnan et al., 2016; Zhou et al., 2020). The first was developed by Zhou and colleagues (2020) (hereafter *Zhou-NS_2020_*), and is a linear Support Vector Machine model trained at discriminating between the sight of hands in pain *vs*. no pain from brain activity within a whole grey matter mask. The second by Krishnan and colleagues (2016) (hereafter *Krishnan_2016_*) is a LASSO multivariate regression, aimed at predicting three different levels of pain in hand images, from principal components of whole brain activity. Both models are described in terms of whole brain weight maps where the value at each coordinate reflects the relative linear contribution of voxelwise activity to the prediction of pain observation. For the purpose of the present study, we took the whole brain maps of each model (available at https://github.com/canlab/Neuroimaging_Pattern_Masks/tree/master/Multivariate_signature_patterns/; files: *“NS_vicarious_pain_pattern_unthresholded.nii”* and *“bmrk4_VPS_unthresholded.nii”*, respectively), and used it to predict the degree of vicarious pain associated with our dataset. We took the first-level *β* parameter estimates associated with each subject/condition, and resampled them to the same resolution of the model weight maps. Vicarious pain was then estimated as the dot-product of the resampled *β*s and the model weights. The higher the resulting value, the higher the correspondance of the first-level images to the pattern expressed by the model. The analysis was carried out with the code available at https://github.com/canlab/Neuroimaging_Pattern_Masks/tree/master/Multivariate_signature_patterns/ (file “apply_all_signatures.m”). The resulting vicarious pain estimates associated with each subject/condition were then fed to the same linear mixed model scheme used for behavioral measures. The only difference from the analysis of behavioral measure lies in the fact that the signature outputs were based on first-level parameter estimate maps, where all repetitions of each subject/condition were collapsed together. As such, we could not model experimental materials (e.g., the specific picture) as random factor. To our knowledge, in the current literature, there is no neural model of vicarious pain response based on verbal material, as such no model-based approach was applied to the “Cognitive and Affective Theory of Mind” task

##### Representational Similarity of Pain

We complemented the above analysis by running a correlation-based Representation Similarity Analysis to identify the presence of any common representation of pain in oneself (from the “Pain Localizer”) and pain in others as perceived through pictures (“Handedness”) and text (“Cognitive and Affective Theory of Mind”). For this purpose, we run first-level GLMs that were identical to those of the standard univariate analysis, except that they were modelled on preprocessed images without normalization and smoothing (Corradi-Dell’Acqua et al., 2011, 2014; Qiao-Tasserit et al., 2018). Following previous studies, we performed a searchlight approach that does not rely on a priori assumptions about informative brain regions, but searches for predictive information throughout the whole brain (Corradi-Dell’Acqua et al., 2011, 2014, 2016; Qiao-Tasserit et al., 2018). For each coordinate in the native brain image, a spherical volume-of-interest was defined around it (5 voxels diameter, 81 voxels total). Then, for each individual subject, we extracted the parameter estimates associated with all conditions of interest within this sphere. Thus, each of the conditions was associated with a unique multivoxel pattern of *β*s in the volume of-interest. These patterns were then correlated one with another, thus resulting in a symmetrical correlation matrix. The correlation coefficients *r* in this matrix were Fisher transformed *z = 0.5 * log_e_[(1 + r)/(1 – r)]* (Corradi-Dell’Acqua et al., 2011, 2014) and then assigned to the center voxel of the sphere.

For the purpose of the present study, we considered the average of the *z*-transformed correlation coefficients obtained when pairing together different pain conditions (*Hot* temperatures, *PF* pictures, *Pain* scenarios & judgments). Such “within-pain” similarity was compared to the average correlation obtained when pairing one pain condition with the painless emotional events from the same paradigm. This resulted in different *z*-maps for each individual, which were then normalized to the MNI template and smoothed using an 8 mm FWHM Gaussian kernel. These maps were then fed to the same flexible factorial routine used for the standard univariate analysis.

## Results

### Cohort 1: University Students

#### Handedness Task

##### Behavioral Responses

We first analyzed data from the Handedness task (Corradi-Dell’Acqua et al., 2011, 2019; Qiao-Tasserit et al., 2018), in which individuals processed hands in Painful *(PF)* or Negative Painless *(PL)* situations, as well as neutral control images *(cPF, cPL)* matched with the previous two categories for visual features, but purged from any emotionally salient characteristics (see Figure 1A). Supplementary Table S1 provides a full description of the behavioral findings. Briefly, participants found more challenging assessing the laterality of emotionally arousing pictures (*PF*, *PL*) compared to their neutral controls *(cPF, cPL)*, as shown by longer Response Times or lower Accuracy (Figure 1B, boxplots higher than 0). However, for painful (*PF*) stimuli, this effect was present only in Controls, and significantly weaker in senior medical students (*Med4*), as revealed by a three-way interaction EMOTIONAL AROUSAL*STIMULI*GROUP associated with the analysis of Response Times. Furthermore, familiarity ratings showed a two-way interaction STIMULI*GROUP, supporting a progressively higher familiarity in medical students (compared to controls) for *PF* stimuli and visually-matched controls *cPF* (Figure 1C), but not for *PL & cPL* pictures.

**Figure 1.**
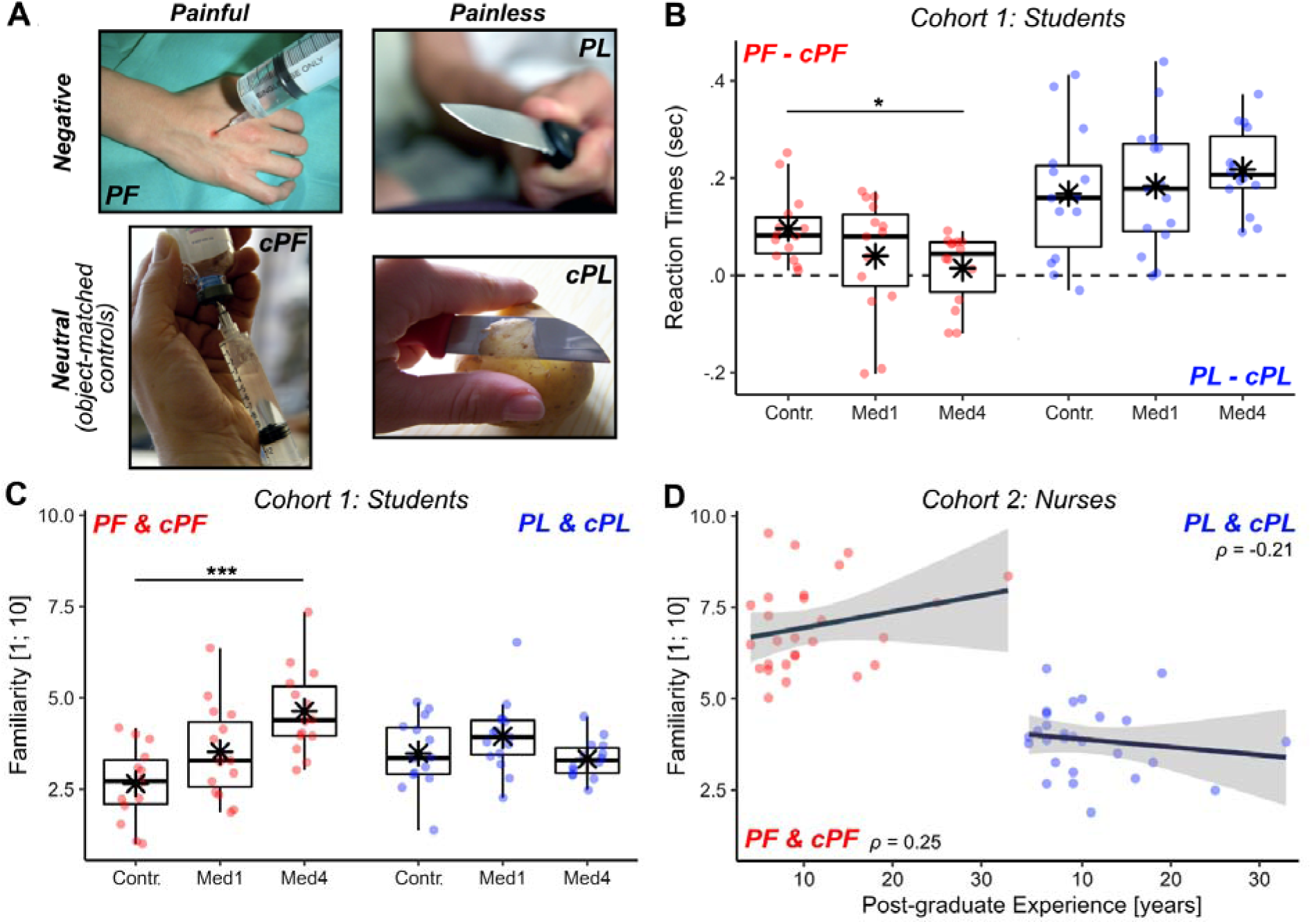
Handedness task. (**A**) Example for each stimulus category. PF: Painful; PL: Negative Painless; cPF & cPL: respective neutral control conditions. (**B-C**) From cohort 1, boxplots displaying mean online Response Times of correct responses and post-scanning Familiarity Ratings. Response Times are displayed as differential seconds between each emotional condition and its neutral control. Familiarity values are displayed as average scores between each conditions and its’ control, and range from 1 (not familiar at all) to 10 (extremely familiar). For each boxplot, the horizontal line represents the median value of the distribution, the star represents the average, the box edges refer to the inter-quartile range, and the whiskers represent the data range within 1.5 of the inter-quartile range. Individual data-points are also displayed color coded, with red dots referring to *PF/cPF* stimuli, and blue dots to *NPL/cNPL* stimuli. *Contr.*: Controls; *Med1 & Med4*: university students enrolled at the first/fourth year of medicine. “***” and “*” refer to significant group differences as tested through linear mixed models (see methods) at *p* < 0.001 and *p* < 0.05 respectively. (**D**) From cohort 2, scatter plot displaying post-scanning Familiarity ratings against post-graduate EXPERIENCE. Each plot is described though a regression line, 95% confidence interval area, color-coded data points, and a Spearman’s rank-correlation coefficient *ρ*. For comparability purposes, Familiarity scores in subplots (C) and (D) are displayed on the same scale.

##### Neural Responses

Supplementary Tables S2-4 provide full details about brain regions recruited during the Handedness task. As expected, a distributed network including the middle (MI) and anterior insula (AI), the middle cingulate cortex (MCC), and the supramarginal/postcentral gyri (SMG/PCG), showed increased activity to pictures of hands in pain (contrast *PF – cPF*) regardless of participants’ healthcare training (Ding et al., 2019; Y. Fan et al., 2011; Jauniaux et al., 2019; Kogler et al., 2020; Lamm et al., 2011; Schurz et al., 2021; Timmers et al., 2018) (Figure 2A, red blobs). Instead, arousing but painless images (*PL – cP L*) activated only AI together with medial cortices around the supplementary motor area (SMA) (Figure 2A, blue blobs).

**Figure 2.**
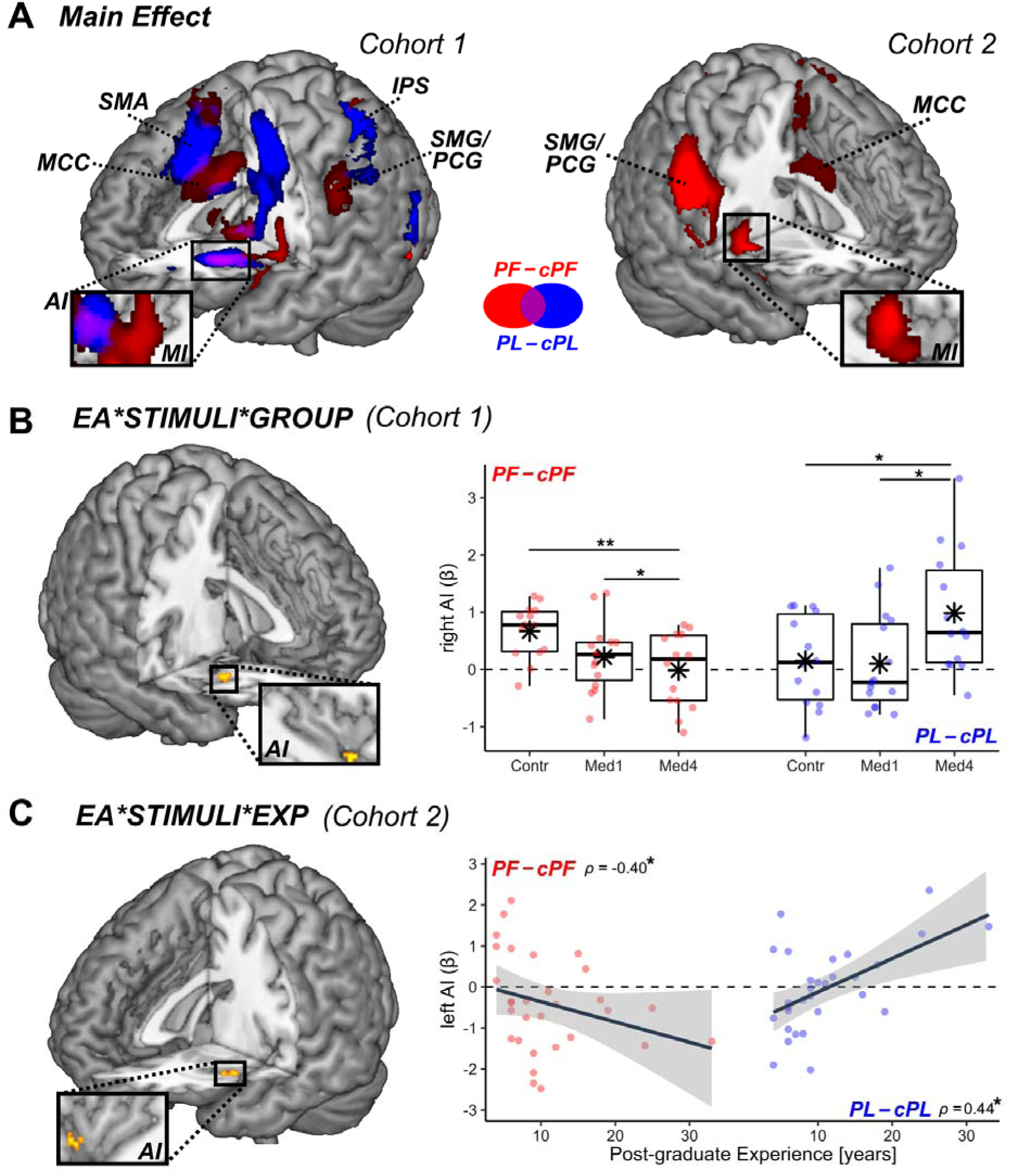
Handedness Task. Surface rendering of brain regions associated with (**A**) the main effect of PF – cPF (red blobs) and PL – cPL (blue blobs) across in both cohorts. SMG: Supramarginal Gyrus; PCG: Postcentral Gyrus; SMA: Supplementary Motor Area; MCC: Middle Cingulate Cortex; MI & AI: Middle & Anterior Insula; PreC: Precentral Gyrus; IFG: Inferior Frontal Gyrus. (**B**) In Cohort 1, regions associated with the three-way interaction between Emotional Arousal (EA), Stimuli, and Group (where “Group” refers to the difference between Control and Med4 students). All activated regions are displayed under TFCE correction for multiple comparisons for the whole brain or mask of interest (see methods). Detailed coordinates are listed in Supplementary Tables S2-4. Activity parameters extracted from the highlighted region are plotted according to groups. Individual data-points are also displayed. Red dots refer to*PF – cPF* activity, whereas blue dots refer to *PL – cPL* activity. “**”, and “*” refer to significant group differences (tested through independent sample t-test) at *p* < 0.01 and *p* < 0.05 respectively. (**C**) In Cohort 2, regions associated with the three-way interaction between EA, Stimuli, and post-graduate Experience. Parameters extracted from the highlighted region are displayed in a scatter plot, describing differential *PF – cPF* or PL – cPL activity against Experience. “*” refers to Spearman’s rank-correlation coefficient *ρ* associated with *p* < 0.05.

We then investigated how these activations were influenced by healthcare experience, and found that right AI was associated with a three-way interaction EMOTIONAL AROUSAL*STIMULI*GROUP. Figure 2B displays the parameter estimates extracted from the latter region, revealing that its response to painful images (*PF – c P*, r*F*ed dots) decreased linearly from *Controls* (the most sensitive group) through to *Med1* and *Med4* (the least sensitive group). Importantly, such decrease was not found for the response to negative painless images (*PL – cPL*, blue dots), which showed even an opposite trend (Fig. 2B).

Overall, our data converge with previous findings that scholarly healthcare experience reduces the reactivity of key brain regions implicated in the processing of others’ pain, such as AI (Cheng et al., 2007, 2017; Dirupo et al., 2021). Critically, this modulation does not reflect a more global hypo-reactivity, as these same regions exhibit enhanced response to other categories of aversive (painless) pictures.

##### Vicarious Pain Signatures

To further characterize these effects of healthcare training on neural responses to observed pain, we applied a model-based approach and fed our dataset to a well-established predictive neural “signature” of vicarious pain defined by brainwide activity. To maximize the comparability with our “Handedness” task, we considered two models derived from previous fMRI work measuring brain responses to the sight of injured limbs: *Zhou-NS_2020_*(Zhou et al., 2020) and *Krishnan_2016_* (Krishnan et al., 2016) (see Methods). Both models aim at predicting the same underlying construct of pain, but do so by relying on the activity of different brain structures: whereas *Zhou-NS_2020_* relies strongly (but not exclusively) on the middle-anterior insula (Zhou et al., 2020) (Figure 3), *Krishnan_2016_* is grounded on a more widespread network involving occipital, parietal, medio-prefrontal, and subcortical structures (Krishnan et al., 2016). When applying these models to our data from the “Handedness” task, we found that both predicted a significant vicarious pain response to the sight of injured hands in Controls (*PF vs. cPF; t* _(13)_ ≥ 2.3 3, *p* ≤ 0.036), demonstrating a reasonable generalizability of these neural patterns to our dataset (at least for individuals with no healthcare training).

**Figure 3.**
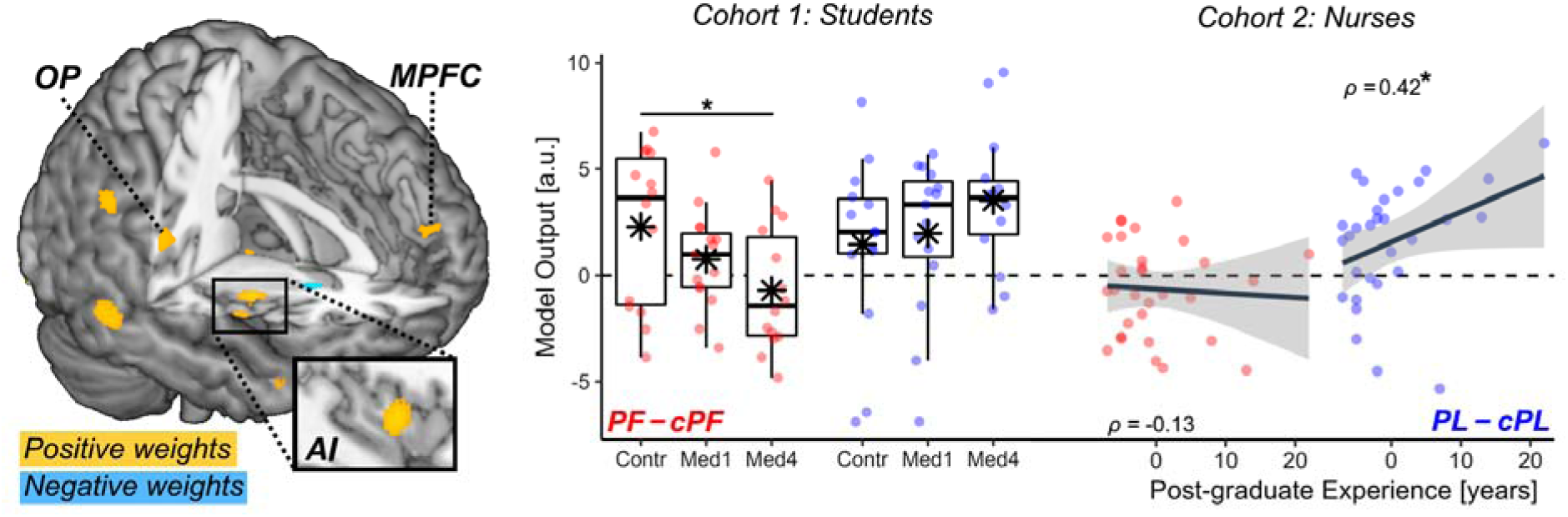
Handedness Task: Vicarious Pain Signatures. Surface brain renderings displaying regions that contributed positively (yellow blobs) and negatively (cyan blobs) to the prediction of vicarious pain from the sight of injured limbs, based on the model of Zhou-NS _2020_. The contribution maps are displayed under a false discovery rate correction of q < 0.05, as provided in the original study (Zhou et al., 2020). The estimated vicarious pain (in arbitrary units [a.u.]) from our data are also displayed. Red dots refer to the differential *PF – cPF* output, whereas blue dots refer to differential *PL – cPL* output. MPFC: Medial Prefrontal Cortex; AI: Anterior Insula; OP: Parietal Operculum. N = sample size. “*” refers to significant group differences (through *t*-statistics) or rank-correlation coefficient *ρ* associated with *p* < 0.05.

We then assessed the degree to which the sensitivity of these models was affected by healthcare education. Full results are reported in Supplementary Table S5. Critically, the output of the model of *Zhou-NS_2020_*was associated with a significant three-way interaction EMOTIONAL AROUSAL*STIMULI*GROUP, similar to our behavioral and whole-brain neuroimaging results. As shown in Figure 3 (red boxplots), the signature fit from this model became progressively less sensitive to injured hands (*PF – cPF*) when moving from *Controls* to *Med1* and *Med4*. This effect did not generalize to the sight of painless images, whose neural signature fit was fairly comparable across groups (*PL – cPL*; Figure 3, blue boxplots). On the other hand, the output of *Krishnan_2016_* model revealed no significant interaction effect, and was reliably sensitive to the sight of injured limbs (*PF – cPF*) in each of the three groups (*t* ≥ 2.18,*p* (1-tailed) ≤ 0.026), with no difference among them.

Finally, as participants from both experiments underwent a brief session in which they received painful hot or painless warm stimulations on their own hand (“Pain Localizer”; see Methods), we also assessed how well these two signature models were sensitive to pain experienced by oneself. Our results replicated previous studies, showing that *Zhou-NS_2020_* output was higher for self-pain (vs. painless thermal stimulation) in all groups (*t* ≥ 4.0 9*p*, ≤ 0.002). However, this was not the case for *Krishnan_2016_*output (| *t*| ≤ 1.39, *p* ≥ 0.187), suggesting that this model might be more sensitive to information independent from first-hand nociception.

Taken together, these data highlight that information about vicarious pain is encoded in a widespread network, captured in different ways by different neural models. This discrepancy reflects the fact that each model relies on different brain structures for their predictions. Importantly, our analysis show that the vicarious pain signature becomes less predictive with increasing medical education, and this effect of healthcare training impacts selectively a model (*Zhou-NS_2020_*) that highlights activity of the insular cortex and encodes information shared with first-hand nociception.

#### Cognitive and Affective Theory of Mind Task

Participants also underwent a previously validated “Cognitive and Affective Theory of Mind” task (Corradi-Dell’Acqua et al., 2014, 2020). In this task, participants read brief story (*scenario epoch*) followed by a question probing for the protagonist’s pain, emotions or beliefs (*judgment epoch*). Supplementary Tables S6 provide a full description of the behavioral data in this task, which revealed no effect of healthcare education on the explicit appraisal of pain, emotions, or beliefs.

For fMRI data, we analyzed brain activity during the scenarios epochs separately from that during the judgment (Aichhorn et al., 2008; Corradi-Dell’Acqua et al., 2014) (see Methods). Whereas judgments require participants to focus on one specific state category, the scenarios could elicit a mix of spontaneous appraisals about different cognitive and affective states. Hence, we took advantage of validation data obtained from an independent population who quantified each narrative in terms of how it triggered inferences about pain, emotions, or beliefs (see methods and Corradi-Dell’Acqua et al., 2014). We applied these scores to the present study in order to identify neural structures, whose activity correlated with each mental state category (Corradi-Dell’Acqua et al., 2014). As described in details in Supplementary Tables S7-S10, our results revealed that text scenarios evoking inferences about pain differentially engaged the MCC, SMG, medial prefrontal cortex (MPFC), and insula, including AI (Bruneau, Dufour, et al., 2012; Bruneau et al., 2015; Bruneau, Pluta, et al., 2012; Corradi-Dell’Acqua et al., 2014, 2020; Jacoby et al., 2016) (Figure 4A, red blobs). On the other hand, scenarios highlighting emotions engaged a partly-similar network involving AI, MPFC and SMG at the border of PCG. In addition, scenarios engaging emotions activated the middle temporal cortex, extending to the temporal pole, as well as the superior parietal cortex (blue blobs). Finally, scenarios referring to beliefs engaged a distinctive network implicating the bilateral temporo-parietal junction, precuneus, and dorsomedial prefrontal cortex, consistent with previous meta-analyses on theory-of-mind and mentalizing (Mar, 2011; Molenberghs et al., 2016; Schurz et al., 2014, 2021; Van Overwalle, 2009). The same networks evoked by pain, emotion, and belief scenarios were also recruited during the judgment epochs (compared to a high-level control condition involving questions about outdated pictures/photos, see methods).

**Figure 4.**
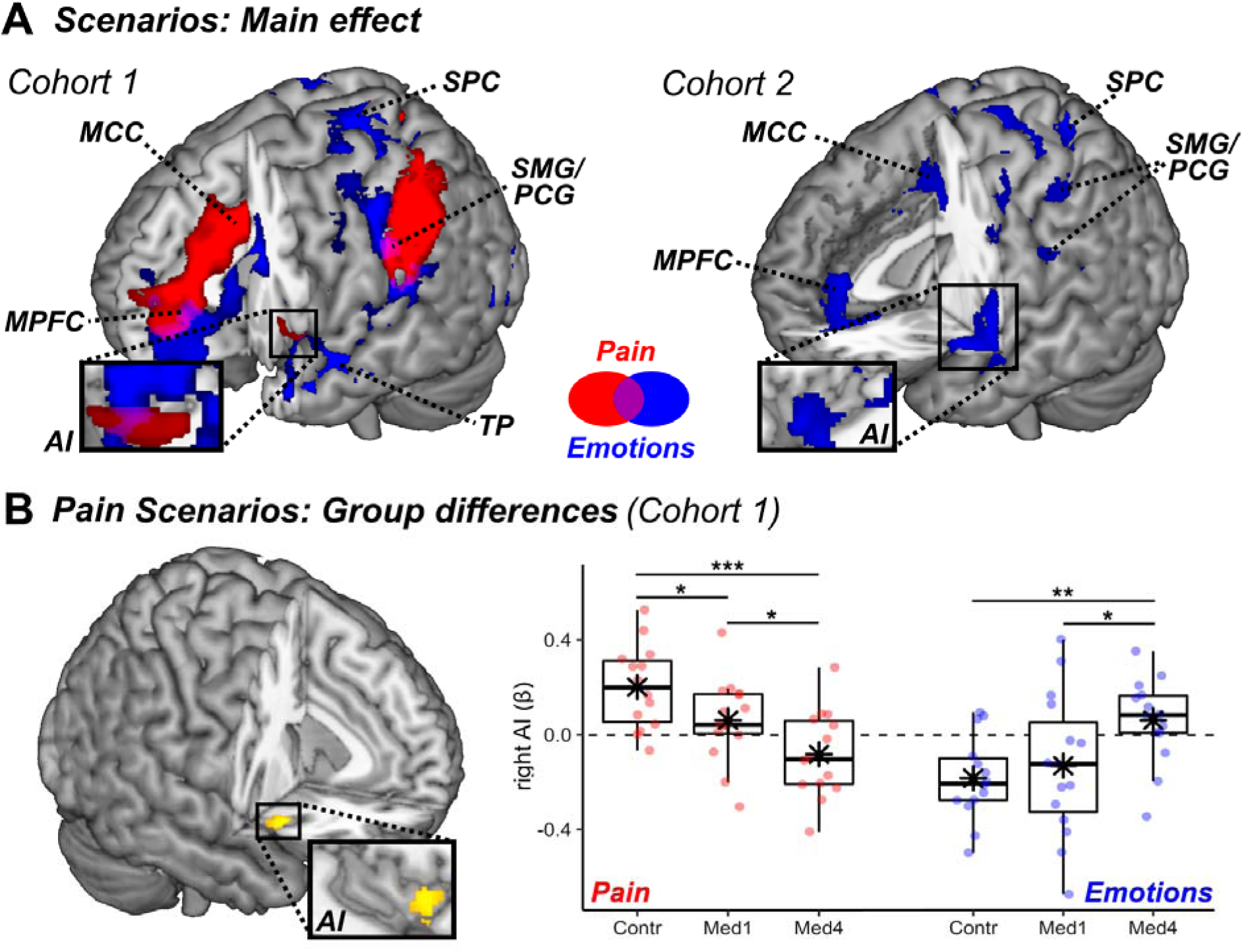
Cognitive and Affective Theory of Mind Task: surface rendering of brain regions associated with (**A**) reading text-based scenarios about individual Pain (red blobs) and Emotions (blue blobs) in both cohorts, (**B**) group differences in Pain scenarios in Cohort 1 (yellow blobs refer to the contrast Control > Med4). All activated regions are displayed under TFCE correction for multiple comparisons for the whole brain or mask of interest (see methods). The parameters extracted from the highlighted region are plotted across groups (with boxplots). Individual data-points are also displayed. Red boxes/lines/dots refer to pain-evoked activity, whereas blue boxes/lines/dots refer to emotion-related activity. Detailed coordinates are listed in Supplementary Tables S8-S9. MPFC: Medial Prefrontal Cortex. SMG: Supramarginal Gyrus; PCG: Postcentral Gyrus; MCC: Middle Cingulate Cortex; SPC: Superior Parietal Cortex; AI: Anterior Insula; TP: Temporal Pole. Contr.: Controls; Med1 & Med4: university students enrolled at the first/fourth year of medicine. N = sample size. “***”, “**”, and “*” refer to significant group differences from independent sample t-tests at p < 0.001, p < 0.01 and p < 0.05 respectively.

We then assessed the role played by healthcare training on these effects by testing for group differences in the recruitment of each of these networks. Results showed that scenarios referring to pain elicited stronger neural responses in the right AI in controls as opposed to nurses. Figure 4B displays the activity parameter estimates from this region who exhibited a progressive decrease of pain-related activity across levels of healthcare training. Interestingly, and similarly to the “Handedness” task, this decrease did not extend to scenarios referring to painless emotions which, instead, showed an opposite trend (Figure 4B, blue dots).

#### Representational Similarity of Pain

Our participants were exposed to pain across different paradigms, some involving hot temperatures delivered to their own body (“Pain Localizer”), and others involving a representation of pain in others, either through images (“Handedness” task) or text (“Cognitive & Affective Theory of Mind” task). To seek for regions disclosing a common representation of pain across these different paradigms, we performed a multi-voxel pattern analysis of neural response allowing us to compare these conditions and test whether any representational similarity between them changed as function of healthcare experience. More specifically, we tested for regions exhibiting the highest pattern similarity whenever two painful conditions were paired together (Hot temperatures; *PF* images of wounded hands; Pain Scenarios & Judgments; Figure 5A, yellow blocks), as opposed to when they were paired with another emotional painless event (*PL* images; Emotion Scenarios/Judgments; Figure 5A, blue blocks). We then estimated pain-specific information by contrasting pattern similarity in “within-pain” vs. “across-affect” pairings (Figure 5A), *via* a whole-brain searchlight analysis. Results revealed a consistent high representational similarity of pain across all groups in the left middle-anterior insula, extending to the left orbitofrontal cortex and bilateral inferior frontal gyrus (IFG) (Figure 5B, left plot; and Supplementary Table S11). No group difference was observed in the voxelwise analysis. Interestingly, however, average similarity scores extracted from this left AI cluster revealed that a reliable “within-pain” vs. “across-affect” difference was present only in the Controls (non-parametric Wilcoxon sign rank test: *Z* = 2.89,*p* = 0.003) but not in the other two groups (*Z* ≤ 1.48, *p* ≥ 0.147). Furthermore, the differential similarity between “within-pain” vs. “across-affect” conditions in AI was stronger in Controls than in the other two groups (Man-Whitney rank sum test: *Z* ≥ 2.07,*p* ≤ 0.038; Figure 5C left subplot), although the effect was not sufficiently strong to exceed the threshold adopted for the voxelwise analysis. Unlike the left AI, the left IFG and MI displayed higher “within-pain” *vs*. “across-affect” similarity in all groups (*Z* ≥ 1.82, *p* (1-tailed) ≤ 0.035), with no significant difference between these conditions (*Z* ≤ 1.24, *p* ≥ 0.214; see Figure 5C, right subplot for one example).

**Figure 5.**
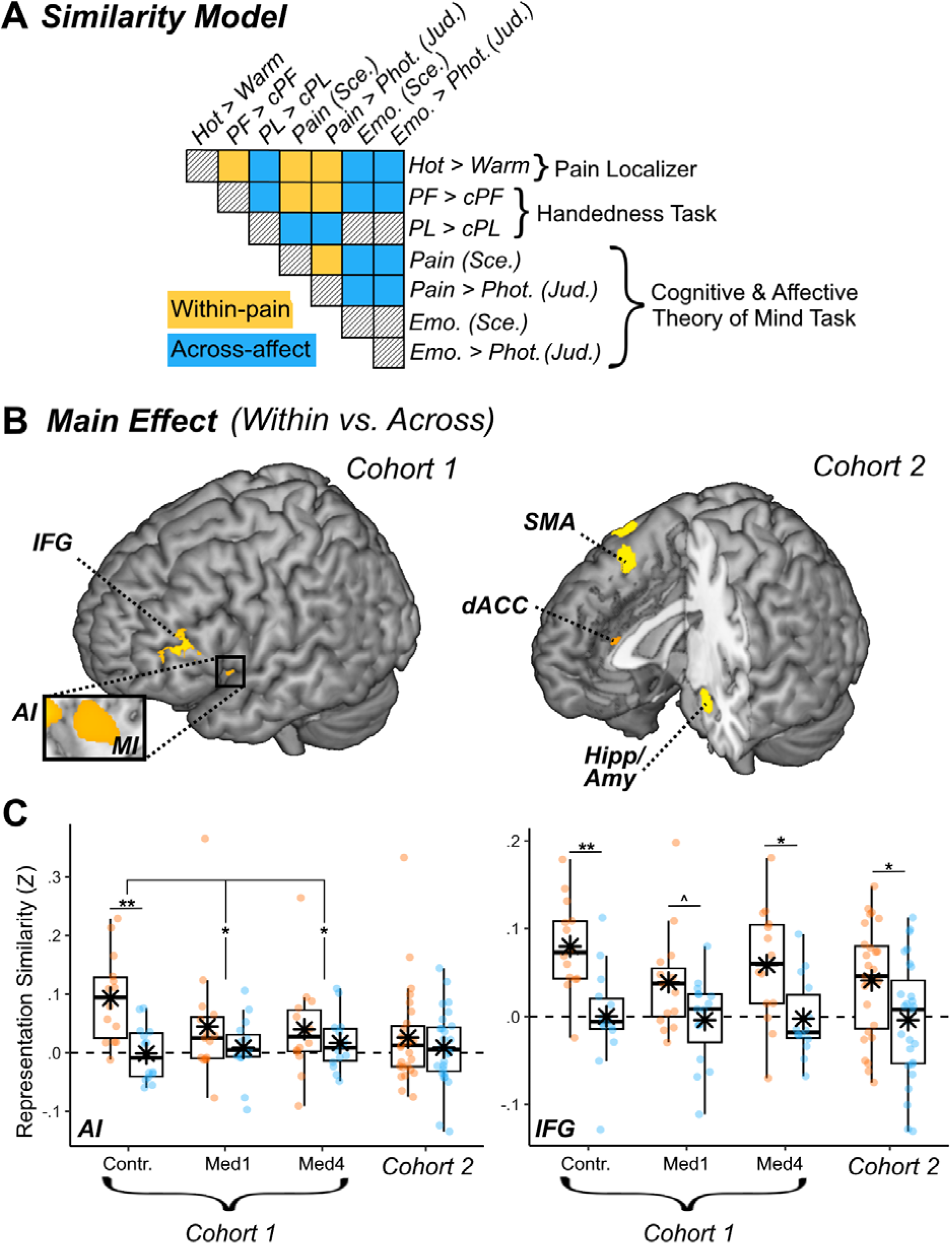
Representational Similarity of Pain. (**A**) Schematic matrix of a representation model testing for pain-specific activity patterns across different tasks in our study. Within the matrix, row and column labels refer to conditions of interest: Hot temperatures from the “Pain Localizer”, Painful (PF) and Negative Painless (PL) images from the “Handedness” task, and Beliefs, Emotions, and Pain Scenarios & Judgments from the “Cognitive and Affective Theory of Mind” task. Each matrix cell indicates the putative representation similarity between two conditions: yellow cells concern pairs of pain conditions (within-pain), blue cells concern pairs of a painful and emotional painless conditions (Across-affect), and black-striped cells refer to pairings of no interest. (**B**) Surface brain rendering of regions displaying significant main effects in their sensitivity to pain-specific information (within-pain vs. across domain). Detailed coordinates are listed in Supplementary Table S12. (**C**) Activity parameters extracted from the highlighted region are plotted across groups and cohorts, with boxplots and individual data. MI & AI: Middle & Anterior Insula; IFG: Inferior Frontal Gyrus; dACC: dorsal Anterior Cingulate cortex; SMA: Supplementary Motor Area; Hipp: Hippocampus; Amy: Amygdala. Contr.: Controls; Med1 & Med4: university students enrolled at the first/fourth year of medicine; “***”, “**”, and “*” refer to significant effects differences from non-parametric rank tests at p < 0.001, p < 0.01 and p < 0.05 respectively.

### Cohort 2: Emergency Nurses

#### Handedness Task

We repeated the same set of analyses on an independent cohort of emergency nurses who underwent a similar protocol. In their case, healthcare training was not only acquired by scholarly education but also consolidated by clinical experience. Figure 2A (right subplot) displays brain regions implicated in processing pain pictures in the Handedness Task (contrast *PF – cPF*) in this group. This analysis confirmed the involvement of a network containing MI, MCC, and the SMG/PCG (Supplementary Table S2). No effect was found in the most anterior portion of the insula. However, and similarly to the cohort 1, AI activity showed a three-way interaction EMOTIONAL AROUSAL*STIMULI*EXPERIENCE. As seen in Figure 2C, this region became less responsive to painful images (*PF – c P*, *F*red dots) as the nurses became more experienced. Furthermore, and similarly to cohort 1, the neural response of AI did not exhibit such decrease to negative painless images (*PL – cPL*, blue dots), but rather showed the opposite trend.

We also analyzed cohort 2 by testing them on the two multivariate “brain signature” models for vicarious pain: *Zhou-NS_2020_* (Zhou et al., 2020) and *Krishnan_2016_* (Krishnan et al., 2016) (see Supplementary Table S5 for full details). When probing for the output of *Zhou-NS_2020_*model, we found that the three-way interaction EMOTIONAL AROUSAL*STIMULI*EXPERIENCE was marginally significant (*t*_(84)_ = -1.76,*p* = 0.082). Figure 3 (left side) displays the model output plotted against post-graduate experience, and reveals that, as in cohort 1, this interaction stems from an effect of experience on the differential sensitivity of the model to *PF* and *PL* images, with a reduced neural signature found for the former relative to the latter. The only discrepancy between the two datasets is that, in cohort 1 the interaction was mainly explainable in terms of lower sensitivity to *PF* (*vs. cPF*), whereas in cohort 2 the effect was driven by a higher response to *PL* (*vs. cPL*) which increased with the number of years of experience. On the other hand, when probing for the output of *Krishnan_2016_* model, we again found a preserved neural signature that was reliably stronger for *PF* (*vs. cPF*) in cohort 2 (*t*_(27)_ = 6.09, *p* < 0.001) like in cohort 1 above, without any modulation of EXPERIENCE (|*t*| ≤ 1.51).

Finally, we tested the sensitivity of the same two neural models using data from the Pain Localizer session. We found that *Zhou-NS_2020_*was reliably sensitive to self-pain (vs. painless temperature) (*t*_(27)_ = 4.69, *p* = 0.002), whereas *Krishnan_2016_* was not (*t*_(27)_ = -0.99, *p* = 0.331).

Overall, our analysis of pain-related responses from the handedness task in emergency nurses provides a conceptual replication of the effects observed for medical students, suggesting that modulations due to scholarly education extend also to experience gathered with clinical exposure. Furthermore, the effects of healthcare training in nurses dominated for responses to seen pain associated with neural activity in AI and partly shared with self-pain.

#### Cognitive and Affective Theory of Mind Task

Results from the Cognitive and Affective Theory of Mind task in cohort 2 confirmed previous findings on beliefs and emotion, with the former recruiting temporo-parietal junction and precuneus (Supplementary Tables S7 & S10), and the latter implicating a widespread network involving AI, MPFC, SMG and temporal cortex (Figure 4A, right plot, Supplementary Tables S8 & S10). The only effect from cohort 1 that was not replicated was the neural modulation associated with the pain condition, as the current sample of nurses showed no suprathreshold effect whatsoever. Finally, no significant modulation was found for the effects associated with EXPERIENCE.

#### Representational Similarity of Pain

As the last step, we repeated the multi-voxel pattern analysis seeking for a common neural representation of pain across tasks in cohort 2. The results are described in Figure 5B (right subplot) and Supplementary Table 11. We observed a high representational similarity of pain at the level of bilateral amygdala and hippocampus, as well as dACC and SMA. No effect of EXPERIENCE was found for these regions. Unlike in cohort 1, our analysis revealed no pain-specific representation at the level of the insula and IFG. Furthermore, when extracting the average similarity scores from a region corresponding to the left AI cluster observed in cohort 1, we found no difference in similarity between “within-pain” vs. “across-affect” pairs even at the uncorrected level (*Z* = 0.05, *p* = 0.962; Figure 5C left subplot). This was not the case of other regions: for instance, the left IFG cluster from cohort 1 displayed higher similarity for the “within-pain” vs. “across-affect” pairings also in cohort 2 (*Z* = 2.47, *p* = 0.013).

### Cohorts comparison

For a sub-part of data that were comparable across groups, we ran a follow-up analysis where we combined the two cohorts together, and examined the effect of healthcare training in terms of a four-level GROUP factor (*Controls*, *Med1*, *Med4*, *Nurses*). Results are fully detailed in Supplementary Tables S12-15 and Figures S1-2, and confirm that, in both the “Handedness” and “Cognitive and Affective Theory of Mind” tasks, female nurses from cohort 2 showed a lower neural response to others’ pain relative to control students. This was observed both for the voxelwise analysis, revealing significant group differences in the right AI, and for the vicarious pain signature from *Zhou-NS_2020_*. This analysis also revealed a significant effect on valence ratings, showing that female nurses rated pictures of injured limbs as less negative than control students. Overall, this follow-up analysis suggests that nurses are less reactive to others’ pain than controls, and more aligned with *Med4*.

## Discussion

We investigated the role of healthcare experience in individuals’ sensitivity to others’ pain, and whether this may extend to other forms of social cognition. We confirmed that healthcare experience decreases the brain response to others’ pain, especially neural activity in AI (Cheng et al., 2007, 2017; Choinière et al., 1990; Davoudi et al., 2008; Dirupo et al., 2021). Importantly, we also demonstrated that such attenuation is not restricted to paradigms using pictures (Cheng et al., 2007, 2017; Decety et al., 2010; Dirupo et al., 2021; Xie et al., 2018), but generalizes to situations where pain was inferred from short narratives. Furthermore, healthcare experience did not influence in the same fashion the neural responses to others’ painless emotional states. For picture-evoked activity, these results were conceptually replicated in a separate dataset from emergency nurses by testing healthcare experience due to professional (rather than scholarly) training. Finally, multivariate pattern analysis revealed that healthcare experience affected specifically a component of the neural representation of other’s pain centered on AI and common with first-hand nociceptive stimulations. Information about others’ pain could still be discriminated from other networks in all participants regardless of their scholarly/professional background.

Previous studies have reported how AI responds to others’ suffering (Ding et al., 2019; Y. Fan et al., 2011; Jauniaux et al., 2019; Kogler et al., 2020; Lamm et al., 2011; Schurz et al., 2021; Timmers et al., 2018), as conveyed by facial expressions (Dirupo et al., 2021; Rütgen et al., 2015, 2021; Wagner et al., 2020; Zhou et al., 2020), photos/videos of injuries (Braboszcz et al., 2017; Cheng et al., 2007, 2017; Corradi-Dell’Acqua et al., 2011, 2019; Karjalainen et al., 2017; Krishnan et al., 2016; Qiao-Tasserit et al., 2018; Zhou et al., 2020), symbolic cues (Berluti et al., 2020; Corradi-Dell’Acqua et al., 2016; López-Solà et al., 2020; O’Connell et al., 2019), or short narratives (Bruneau, Dufour, et al., 2012; Bruneau et al., 2015; Bruneau, Pluta, et al., 2012; Corradi-Dell’Acqua et al., 2014, 2020; Jacoby et al., 2016). Yet the functional interpretation of these neural responses to pain remains debated. One proposal is that a large part of this activity might encode supraordinal features, such as unpleasantness, arousal, or salience (Corradi-Dell’Acqua et al., 2011, 2016; Iannetti & Mouraux, 2010; Mouraux et al., 2011).

Another view, based on studies carefully controlling for supraordinal confounders, assumes that both self and vicarious pain are represented in the insular cortex in a pain-selective manner, perhaps parallel to other supraordinal signals (Corradi-Dell’Acqua et al., 2016; Horing et al., 2019; Liang et al., 2019; Rütgen et al., 2021; Sharvit et al., 2018, 2020). Our results converge with, but also extend, the latter notion by showing that healthcare experience changes AI responses to others’ suffering in a pain-specific fashion. In contrast, AI still responded to painless emotional pictures and narratives in caregiver groups, with this reactivity impacted by healthcare experience in a different way, characterized by unchanged or even increased (rather than decreased) activation in more senior students/nurses.

Our results therefore fit with the idea that healthcare experience modifies neuronal populations sensitive specifically to pain. Alternatively, it is still theoretically possible that healthcare experience affects neuronal populations in AI that code for broader properties like unpleasantness, salience, or avoidance. In this case, however, the influence of healthcare experience would still be dependent of the nature of the state observed in others (pain vs. other affective states), and cannot be explained by a general hypo-reactivity or inhibition. For instance, it is conceivable that healthcare experience could modulate the “access” of vicarious pain cues to neurons coding for these broader signals, by regulating the likelihood for a specific type of stimuli to engage this region. Indeed, seminal models suggest that AI is also part of the so-called “salience network”, responding to events sufficiently relevant to capture attention and motivate behavioral adjustment (Iannetti & Mouraux, 2010; Mouraux et al., 2011; Uddin, 2015). Our data might arguably be compatible with this interpretation, as professional exposure to injuries, cuts, or burns would make these categories less novel or unusual, and therefore less likely to trigger neurons responding to salience, at the advantage of other events outside this domain of expertise.

We also performed comprehensive multivariate pattern analyses to examine the similarity between the neural response to pain information arising from different cues, and relative to other forms of affect. By examining two previously established and independently defined neural signatures of vicarious pain, based on whole-brain activity patterns, we found that healthcare experience impacted exclusively the proficiency of a model strongly grounded on AI and sensitive also to self-pain (Zhou et al., 2020). Instead, when using another neural model, derived from activity in more widespread regions and independent from first-hand nociception (Krishnan et al., 2016), we could still correctly predict responses to pain pictures in all groups to the same degree, regardless of healthcare training. These results were complemented by a representational similarity analysis, which revealed that healthcare experience disrupts the neural overlap between pain signals in AI across our different paradigms (temperatures, pictures, and verbal scenarios), but not that of other regions (e.g., IFG). Furthermore, even in experienced healthcare providers, pain-specific information could be inferred from the neural activity of non-insular structures, such as dACC, SMA or amygdala/hippocampus, consistently with previous studies using similar approaches (Corradi-Dell’Acqua et al., 2016; O’Connell et al., 2019; Wagner et al., 2020). Taken together, these results further support the role of AI in the representation of others’ pain through the recruitment of neuronal populations that are also involved in processing pain in one’s own body (Braboszcz et al., 2017; Corradi-Dell’Acqua et al., 2011, 2016; Qiao-Tasserit et al., 2018). Critically, individuals with longer medical training do not appear to process others’ pain through this pathway, but still rely on information encoded in other brain networks.

Representational similarity between responses to one’s and others’ pain in regions such AI has been often interpreted as a neural substrate for the ability to embody, and empathize with, others’ states (Bastiaansen et al., 2009; Bernhardt & Singer, 2012). In the current study we did not measure individual empathy traits to corroborate this claim. However, in the Handedness task, we did collect implicit (RTs for laterality judgments) and explicit (valence ratings on pictures) measures reflecting individual sensitivity to the affective content of painful stimuli, and found these were indeed influenced by healthcare experience, similarly to brain activity (see Figure 1B & Supplementary Figure S1). In this perspective, our results converge with a wide neuroscience literature suggesting how AI activity in these tasks directly relates to the subjective affective component of responses to one’s and others’ pain (Lamm et al., 2011; Singer et al., 2004; Singer & Lamm, 2009). Hence, the effect of healthcare experience in AI activity could be interpreted in terms of enhanced regulation of “automatic” affective/empathetic responses, allowing repetitive interactions with patients without contagion from their suffering (Gleichgerrcht & Decety, 2014; Vaes & Muratore, 2013).

However, we advise caution in interpreting the modulations observed here in terms of changes in empathy. Empathy is a multidimensional process and neuroimaging results from paradigms like ours do not systematically correlate with scores from validated questionnaires (see, Lamm et al., 2011, for meta-analytic evidence). Furthermore, in contrast with the consistent neuroimaging results (Cheng et al., 2007, 2017; Dirupo et al., 2021; Jackson et al., 2017), behavioral research investigating how healthcare experience impacts on trait empathy has led to mixed results. Some studies suggested that medical students and healthcare providers have lower empathy (Bellini & Shea, 2005; Hojat et al., 2009; Neumann et al., 2011; K. E. Smith et al., 2017), and others report no change (Cameron & Inzlicht, 2020) or even an increase (Chen et al., 2022; Handford et al., 2013; Kataoka et al., 2009; K. E. Smith et al., 2017). This could reflect heterogeneity in definitions and measures of empathy, which vary extensively across studies.

For these reasons, we favor a more parsimonious explanation, suggesting that healthcare experience impacts negatively the affective and embodied processes used to appraise others’ pain, whereby observed somatic/emotional experiences are simulated on oneself. This does not imply a broader reduction in sensitivity to other kinds of painless emotional states, in accordance with the multidimensional psychological construct of empathy captured by questionnaires and other measures.

Finally, a wealth of studies have repeatedly associated empathy traits and the ability to simulate people’s pain (toghether with associated neural activity in regions like AI and dACC) with a wide range of prosocial behaviours, such as altruistic endowment (Ma et al., 2011; Tomova et al., 2017), costly pain relief (Hartmann et al., 2022; Hein et al., 2010), and even organ donation (O’Connell et al., 2019). In this perspective, it could be argued that the effects observed in the present research relate to decreasing prosocial attitudes with increasing healthcare experience, including a well-known tendency to underestimate and undertreat the pain expressed by patients (Choinière et al., 1990; Davoudi et al., 2008; Ruben et al., 2015, 2018; Rupp & Delaney, 2004). We believe this to be only partially true. Although it is plausible that experience-derived changes in AI might lead to decreased pain assessment, this was not directly probed here as we measured only implicit/spontaneous brain reactions to aversive events. However, previous research found that medical education reduced the recruitment of AI and dACC during explicit pain ratings of facial expressions (Dirupo et al., 2021). This being said, however, prosocial behaviours in healthcare environment do not necessarily translate into higher tendency to share and relieve patients’ pain. As pain treatments often involve pharmacological means, physicians and nurses are also concerned with the potential negative contraindications and unwanted consequences of potent painkillers (Bennett & Carr, 2002; Bertrand et al., 2021). Hence, healthcare providers need to balance the deontological goal of alleviating the patients’ *current* suffering with the opposed deontological directive of preventing *future* adverse effects, a conflict that might in some cases lead to withholding analgesia for the patient’s best interest (Corradi-Dell’Acqua et al., 2019). In this perspective, we previously found that AI activity evoked by errors and negative feedback explained nurses’ tendency to hold back drug administration for pain treatment in fear of potential contraindications to painkillers (Corradi-Dell’Acqua et al., 2019). While this accords with the notion that AI activity contributes to prosocial decision making and varies with years of healthcare experience, its recruitment and relationship to actual behavior and medical care in real life are likely to depend on several other factors.

There are few important limitations in our study that should be underlined. First, we investigated the role of healthcare experience by aggregating data from different populations with different levels of experience. As such, our study shares the weaknesses of cross-sectional investigations (Wang & Cheng, 2020), as the role of healthcare experience was not tested longitudinally in the same population. Second, although our sample was sizeable (N = 73), it was split into smaller subgroups (N = from 14 to 30) in order to compare experience effects. As such, not all results from our different paradigms were replicated across the two cohorts, a variability that could relate to low sensitivity, or individual heterogeneity. Third, some stimuli in the handedness task involved medical procedures (injections, surgery) that might appear painful only to lay observers, raising the question as to whether our results truly reflect decreased sensitivity to others’ suffering or rather increased knowledge of the real (e.g. limited) nociceptive impact of these situation. Given that healthcare experience also influenced neural response to pain narratives from the Cognitive and Affective Theory of mind task, which concerned only mundane events unrelated to medical procedures, we are confident that our findings reflect more general changes in pain processing beyond purely medical conditions.

Finally, although the two cohorts underwent almost identical paradigms, they were nonetheless engaged in independent experiments, each with their own idiosyncratic properties. For instance, data from cohort 2 (nurses) were obtained in a larger project involving a wide range of tasks (Corradi-Dell’Acqua et al., 2019), each paradigm was administered in a shorter version relative to cohort 1. Despite this, results from the “Handedness” task revealed a remarkable convergence of the effects of scholarly (cohort 1) and professional (cohort 2) healthcare experience on reactivity to others’ pain, suggesting that they arose over and above all differences between the two datasets. Furthermore, differences between cohorts did not prevent comparing nurses and students for a portion of the data that was matched between the two groups, while also controlling for age. This auxiliary analysis further pointed to reliable effects of experience independent of gender or seniority, although some factors could not be fully equated such as the duration of scanning sessions or fMRI sequence parameters. However, we believe that the impact of these confounds was negligible.

In sum, our study extends previous investigations on the role of healthcare experience in pain processing and social cognition in several ways. First, it shows how healthcare experience influences negatively neural reactivity in AI to others’ pain, both from visual and text information. Second, it shows how this effect is specific for pain, and dissociates from other forms of social cognition, such as painless affect or theory-of-mind abilities. Third, it shows how the neural signature of vicarious pain modified by healthcare experience impacts prevalently the representation in AI, shared with first-hand nociception. Fourth, it demonstrates that, in contrast, information about others’ pain encoded in other brain structures is unaffected by healthcare experience, such that it can be reliably used by predictive multivariate models to detect the sight of injuries. Overall, healthcare experience may result in lower propensity to process others’ suffering as one’s own, accompanied with lower neural reactivity of areas such as AI. These results may contribute to better understand how pain is evaluated and often underestimated in real-life clinical settings (Choinière et al., 1990; Davoudi et al., 2008; Kappesser et al., 2006; Teske et al., 1983).

## Acknowledgments

This study was conducted at the Brain and Behaviour Laboratory (BBL) at the University of Geneva and benefited from the support of the BBL technical staff.

## Authors Contribution

CCD, OH, and PV conceived the design. CCD & CH acquired the data from cohort 1. CCD & GS acquired the data from cohort 2. CCD analyzed the data. OH, PV contributed to the interpretation of the results. CCD drafted the manuscript. All authors revised critically the manuscript.

## Funding

CCD is supported by the Swiss National Science Foundation (SNSF) grant nos. PP00O1_157424, PP00P1_183715 and 320030_182589. PV is supported by the SNSF grant no 32003B_138413.

## Competing Interests

The authors declare they have no competing interests.

## Data and code availability statement

Stimuli, de-identified data and code for the experiment/analysis are available under the Open Science Framework: https://osf.io/8bjmq/

## Notes

https://osf.io/8bjmq/

### Competing Interest Statement

The authors have declared no competing interest.

### Summary of Updates

Minor changes were carried out following peer review

