## Supplements for "Healthcare experience affects pain-specific responses to others’ suffering in the anterior insula"

### Supplementary Information

##### Supplementary Table S1

Handedness Task. Results from (Generalized) Linear Mixed Model analysis with EMOTIONAL AROUSAL (*Neutral, Negative*) and STIMULI (*Painful, Painless*) as within-subjects factor. In Cohort 1, GROUP (*Controls Med1, Med4*) was modeled as between-subjects factor. In Cohort 2, postgraduate EXPERIENCE was modeled as continuous predictor. The *lmer-syntax* of the tested model is the following:

Cohort 1 – *Dep. Var. ~ EA*STIMULI*GROUP + (EA*STIMULI|Subjects) + (GROUP|Materials)*.

Cohort 2 – *Dep. Var. ~ EA*STIMULI*EXP + (EA*STIMULI|Subjects) + (EXP|Materials)*.

The analysis was run on post-experimental ratings of Familiarity, Pain, Arousal and Valence (data from 3 subjects is missing from Cohort 2) as well as on on-line Accuracy and Reaction Times of correct responses. For each dependent variable (displayed vertically), and for each effect of interest (horizontally), the table reports the *Z*/*t*-value associated with the model parameters. Significant effects are highlighted.

|  | ***Accuracy*** | ***RTs*** *[sec]* | ***Arousal*** *[1; 10]* | ***Valence*** *[-4; 4]* | ***Pain*** *[1; 10]* | ***Familiarity*** *[1; 10]* |
| --- | --- | --- | --- | --- | --- | --- |
| ***COHORT 1*** |  |  |  |  |  |  |
| *EMOTIONAL AROUSAL* [EA] | *Z* = **-2.51^‡^** | *t*_(171)_ = **2.23^‡^** | *t*_(68)_ = **13.11^*^** | *t*_(103)_ = -**12.46^*^** | *t*_(76)_ = -**17.74^*^** | *t*_(68)_ = -**6.69^*^** |
| *STIMULI* | *Z* = -0.18 | *t*_(158)_ = 1.45 | *t*_(105)_ = 1.56 | *t*_(117)_ = **4.10^*^** | *t*_(164)_ = -**3.06^†^** | *t*_(75)_ = -1.32 |
| *GROUP* [GR]*: Contr. vs. Med1* | *Z* = 1.41 | *t*_(40)_ = 0.30 | *t*_(40)_ = 0.98 | *t*_(41)_ = 1.68 | *t*_(40)_ = 0.50 | *t*_(40)_ = 1.36 |
| *GROUP* [GR]*: Contr. vs. Med4* | *Z* = **2.11^‡^** | *t*_(40)_ = 0.14 | *t*_(43)_ = 0.27 | *t*_(46)_ = -0.12 | *t*_(41)_ = 1.46 | *t*_(41)_ = 3.68 |
| *EA*STIMULI* | *Z* = -1.47 | *t*_(170)_ = 0.42 | *t*_(80)_ = **-4.91^*^** | *t*_(120)_ = -0.29 | *t*_(80)_ = **-9.51^*^** | *t*_(122)_ = 1.03 |
| *EA*GR: Contr. vs. Med1* | *Z* = 0.73 | *t*_(112)_ = -1.38 | *t*_(40)_ = -1.77 | *t*_(41)_ = 0.02 | *t*_(40)_ = -1.15 | *t*_(40)_ = 0.29 |
| *EA*GR: Contr. vs. Med4* | *Z* = -0.13 | *t*_(144)_ = -**2.25^‡^** | *t*_(45)_ = -1.42 | *t*_(50)_ = 1.66 | *t*_(41)_ = -0.34 | *t*_(44)_ = -1.03 |
| *STIMULI*GR: Contr. vs. Med1* | *Z* = -0.68 | *t*_(76)_ = 1.57 | *t*_(41)_ = 0.82 | *t*_(41)_ = -1.26 | *t*_(41)_ = -0.07 | *t*_(41)_ = -0.52 |
| *STIMULI*GR: Contr. vs. Med4* | *Z* = -0.80 | *t*_(76)_ = -0.78 | *t*_(52)_ = 0.85 | *t*_(52)_ = -1.35 | *t*_(44)_ = -0.95 | *t*_(45)_ = **-4.44^*^** |
| *EA*STIM*GR: Contr. vs. Med1* | *Z* = 0.06 | *t*_(98)_ = 1.20 | *t*_(41)_ = 0.97 | *t*_(41)_ = -0.63 | *t*_(40)_ = 0.83 | *t*_(41)_ = 0.53 |
| *EA*STIM*GR: Contr. vs. Med4* | *Z* = 0.07 | *t*_(111)_ = **2.29^‡^** | *t*_(47)_ = 1.08 | *t*_(53)_ = -0.67 | *t*_(41)_ = -0.20 | *t*_(53)_ = 1.12 |
| ***COHORT 2*** |  |  |  |  |  |  |
| *EMOTIONAL AROUSAL* [EA] | *Z* = **-2.92^†^** | *t*_(106)_ = **2.05^‡^** | *t*_(48)_ = **8.60^*^** | *t*_(57)_ = -**5.91^*^** | *t*_(36)_ = **13.07^*^** | *t*_(70)_ = **-5.35^*^** |
| *STIMULI* | *Z* = -1.03 | *t*_(107)_ = 0.33 | *t*_(78)_ = **3.94^*^** | *t*_(121)_ = -0.39 | *t*_(68)_ = **-2.32^‡^** | *t*_(109)_ = **-7.82^‡^** |
| *EXPERIENCE* [EXP] | *Z* = 1.27 | *t*_(28)_ = 0.94 | *t*_(26)_ = -1.64 | *t*_(25)_ = -0.89 | *t*_(25)_ = -1.05 | *t*_(26)_ = 1.53 |
| *EA*STIMULI* | *Z* = 0.60 | *t*_(112)_ = -0.35 | *t*_(59)_ = -0.62 | *t*_(97)_ = **-2.79^*^** | *t*_(32)_ = **-5.71^*^** | *t*_(80)_ = 0.61 |
| *EA*EXP* | *Z* = -1.18 | *t*_(28)_ = 0.62 | *t*_(25)_ = -0.65 | *t*_(25)_ = 0.26 | *t*_(25)_ = 0.52 | *t*_(25)_ = 0.32 |
| *STIMULI*EXP* | *Z* = -1.62 | *t*_(40)_ = 0.07 | *t*_(26)_ = -0.18 | *t*_(25)_ = 0.14 | *t*_(25)_ = 0.88 | *t*_(26)_ = -1.00 |
| *EA*STIMULI*EXP* | *Z* = 1.43 | *t*_(32)_ = 0.32 | *t*_(25)_ = 0.73 | *t*_(25)_ = 0.25 | *t*_(25)_ = -0.72 | *t*_(25)_ = -0.72 |
| **p < 0.001; †p < 0.01; ‡p < 0.05* | | | | | | |

##### Supplementary Table S2

Handedness Task. Regions displaying differential activity for the contrast *PF > cPF*. L and R refer to the left and right hemisphere, respectively. M refers to medial activations. Activations are displayed under Threshold-Free Cluster Enhancement (TFCE) approach. Cluster size is displayed in terms of consecutive voxels.

|  | ***SIDE*** | ***Coordinates*** | | | ***TFCE*** | ***Cluster Size*** |
| --- | --- | --- | --- | --- | --- | --- |
|  |  | ***x*** | ***y*** | ***z*** |  |  |
| ***Cohort 1, Main Effect: PF > cPF*** | | | | | | |
| Anterior Insula [AI] | R | 34 | 22 | 8 | 1776.57^†^ | 33261 |
| Anterior Insula [AI] | L | -32 | 24 | 2 | 2039.35^*^ |  |
| Posterior Insula [PI] | R | 40 | -2 | -4 | 3474.35^*^ |  |
| Posterior Insula [PI] | L | -40 | 2 | -8 | 2029.61^*^ |  |
| Amygdala | R | 22 | -4 | -16 | 3549.12^*^ |  |
| Amygdala | L | -22 | -4 | -20 | 3273.88^*^ |  |
| Inferior Frontal Gyrus [IFG] | R | 44 | 38 | 6 | 2040.99^*^ |  |
| Inferior Frontal Gyrus [IFG] | L | -50 | 36 | 12 | 866.60^†^ |  |
| Precentral Gyrus | R | 50 | 8 | 20 | 1038.90^†^ |  |
| Precentral Gyrus | L | -46 | 2 | 26 | 1468.67^†^ |  |
| Supramarginal/Postcentral Gyrus [SMG/PCG] | R | 62 | -20 | 36 | 4141.25^*^ |  |
| Supramarginal/Postcentral Gyrus [SMG/PCG] | L | -54 | 24 | 36 | 1503.63^†^ |  |
| Intraparietal Sulcus | R | 24 | -62 | 52 | 3921.23^*^ |  |
| Intraparietal Sulcus | L | -24 | -58 | 54 | 3264.51^*^ |  |
| Inferior Temporal Gyrus | R | 48 | -62 | -10 | 11847.33^*^ |  |
| Inferior Temporal Gyrus | L | -46 | -64 | -10 | 10987.75^*^ |  |
| Middle Occipital Gyrus | R | 34 | -82 | 16 | 11166.37^*^ |  |
| Middle Occipital Gyrus | L | -30 | -88 | 14 | 8731.41^*^ |  |
| Fusiform Gyrus | R | 30 | -56 | -12 | 11616.96^*^ |  |
| Fusiform Gyrus | L | -28 | -54 | -14 | 13705.47^*^ |  |
| Occipital Pole | R | 18 | -94 | -2 | 15702.10^*^ |  |
| Occipital Pole | L | -14 | -90 | -10 | 11892.50^*^ |  |
| Periaqueductal Gray/Midbrain | M | -8 | -24 | -8 | 2160.31^*^ |  |
| dorsal Anterior Cingulate Cortex [dACC] | M | 2 | 4 | 30 | 1596.42^†^ | 1035 |
| Supplementary Motor Area | M | 10 | 14 | 70 | 799.50^†^ | 558 |
| ***Cohort 2, Main Effect: PF > cPF*** | | | | | | |
| Middle Insula [MI] | R | 42 | -2 | 4 | 1664.05^†^ | 595 |
| Precentral Gyrus | R | 50 | 8 | 24 | 606.15^‡^ | 271 |
| Middle Insula [MI] | L | -40 | -6 | 2 | 1392.84^‡^ | 2023 |
| Precentral Gyrus | L | -44 | 4 | 22 | 1010.90^‡^ |  |
| Amygdala | L | -24 | -4 | -22 | 717.93^‡^ |  |
| Hippocampus | L | -22 | -32 | -10 | 665.56^‡^ |  |
| Thalamus | L | -20 | -16 | 4 | 464.08^‡^ | 1 |
| Middle Frontal Gyrus | R | 46 | 40 | 10 | 536.18^‡^ | 39 |
| Middle Orbital Gyrus | L | -24 | 34 | -14 | 465.36^‡^ | 1 |
| Supramarginal/Postcentral Gyrus [SMG/PCG] | L | -58 | -24 | 24 | 4471.27^*^ | 5585 |
| Intraparietal Sulcus | L | -22 | -64 | 48 | 1202.95^†^ |  |
| Middle Cingulate Cortex | M | -2 | 8 | 30 | 651.15^‡^ | 1153 |
| Posterior Cingulate Cortex | M | 2 | -34 | 28 | 539.49^‡^ |  |
| Supplementary Motor Area | M | -10 | -2 | 42 | 590.92^‡^ |  |
| Inferior Temporal Gyrus | R | 52 | -52 | -14 | 2961.58^*^ | 13983 |
| Middle Occipital Gyrus | R | 32 | -82 | 14 | 8122.82^*^ |  |
| Fusiform Gyrus | R | 28 | -48 | -14 | 8517.36^*^ |  |
| Occipital Pole | R | 12 | -86 | -2 | 8857.29^*^ |  |
| Intraparietal Sulcus | R | 26 | -58 | 50 | 2149.88^*^ |  |
| Supramarginal/Postcentral Gyrus [SMG/PCG] | R | 62 | -22 | 38 | 3227.61^*^ | 13983 |
| Inferior Temporal Gyrus | L | -48 | -60 | -10 | 2865.09 |  |
| Middle Occipital Gyrus | L | -26 | -86 | 10 | 8059.38^*^ |  |
| Fusiform Gyrus | L | -28 | -48 | -14 | 7787.88^*^ |  |
| Occipital Pole | L | -18 | -80 | -8 | 11406.56^*^ |  |
| * *p* < 0.001; † *p* < 0.01; ‡ *p* < 0.05 FWE corrected for multiple comparisons for the whole brain.  § *p* < 0.05 Small Volume Corrected for multiple comparisons. | | | | | | |

##### Supplementary Table S3

Handedness Task. Regions displaying differential activity for the contrast *PL > cPL*.

|  | ***SIDE*** | ***Coordinates*** | | | ***TFCE*** | ***Cluster size*** |
| --- | --- | --- | --- | --- | --- | --- |
|  |  | ***x*** | ***y*** | ***z*** |  |  |
| ***Cohort 1, Main Effect: PL > cPL*** | | | | | | |
| Anterior Insula [AI] | R | 44 | 18 | 4 | 2249.70^*^ | 47910 |
| Anterior Insula [AI] | L | -32 | 28 | -4 | 2993.88^*^ |  |
| Dorsolateral Prefrontal Cortex [DLPFC] | R | 28 | 10 | 50 | 3099.96^*^ |  |
| Dorsolateral Prefrontal Cortex [DLPFC] | L | -26 | -2 | 48 | 4035.27^*^ |  |
| Superior Parietal Cortex | R | 42 | -40 | 48 | 3652.86^*^ |  |
| Superior Parietal Cortex | L | -32 | -44 | 42 | 6027.10^*^ |  |
| Middle/Inferior Occipital Gyrus | R | 50 | -66 | 12 | 3210.74^*^ |  |
| Middle/Inferior Occipital Gyrus | L | -40 | -70 | 14 | 3929.21^*^ |  |
| Cerebellum | L | -22 | -54 | -28 | 1194.81^†^ |  |
| Supplementary Motor Area [SMA] | M | -6 | 16 | 44 | 3922.09^*^ |  |
| posterior Middle Cingulate Cortex | M | -4 | -22 | 30 | 2918.73^*^ |  |
| Precuneus | M | -8 | -64 | 42 | 4681.45^*^ |  |
| Periaqueductal Gray/Midbrain | M | -4 | -26 | -14 | 1830.27^‡^ |  |
| dorsal Anterior Cingulate Cortex [dACC] | M | 6 | 24 | 36 | 167.50^§^ | 101 |
| Cerebellum | R | 32 | -62 | -32 | 1033.48^‡^ | 190 |
| ***Cohort 1, (PL > cPL)_Med4_* > *(PL > cPL)_Med1_*** | | | | | | |
| Thalamus | R | 8 | -2 | -2 | 879.36 | 37 |
| Caudate | R | 12 | 12 | 2 | 783.29^‡^ | 14 |
| ***Cohort 2, Main Effect: PL > cPL*** | | | | | | |
| Postcentral Gyrus | L | -40 | -26 | 58 | 2176.32^*^ | 1337 |
| ***Cohort 2, (PL > cPL)*Post-Graduate Experience*** | | | | | | |
| Anterior Insula [AI] | L | -40 | -18 | -12 | 139.03^§^ | 39 |
| * *p* < 0.001; † *p* < 0.01; ‡ *p* < 0.05 FWE corrected for multiple comparisons for the whole brain.  § *p* < 0.05 Small Volume Corrected for multiple comparisons. | | | | | | |

##### Supplementary Table S4

Handedness Task. Regions displaying suprathreshold activity for the interaction (*PF > cPF) >* (*PL > cPL)*.

|  | ***SIDE*** | ***Coordinates*** | | | ***TFCE*** | ***Cluster size*** |
| --- | --- | --- | --- | --- | --- | --- |
|  |  | ***x*** | ***y*** | ***z*** |  |  |
| ***Cohort 1, Main Effect: (PF > cPF) > (PL > cPL)*** | | | | | | |
| Posterior Insula [PI] | R | 42 | -2 | -4 | 1494.25^†^ | 14931 |
| Posterior Insula [PI] | L | -40 | -2 | -6 | 1585.82^†^ |  |
| Amygdala | R | 22 | -4 | -18 | 1391.57^†^ |  |
| Amygdala | L | -22 | -4 | -22 | 2165.63^*^ |  |
| Fusiform Gyrus | R | 28 | -48 | -20 | 7901.24^*^ |  |
| Occipital Pole | R | 14 | -94 | 8 | 10922.80^*^ |  |
| Inferior Temporal/Occipital Gyrus | R | 52 | -60 | -14 | 4158.35^*^ |  |
| Fusiform Gyrus | L | -28 | -56 | -14 | 8248.71^*^ |  |
| Occipital Pole | L | -20 | -90 | -14 | 8508.68^*^ |  |
| Inferior Temporal/Occipital Gyrus | L | -44 | -64 | -8 | 3627.07^*^ |  |
| Posterior Orbital Gyrus | R | 30 | 32 | -20 | 606.69^‡^ | 32 |
| Supramarginal/Postcentral Gyrus [SMG/PCG] | R | 62 | -20 | 28 | 1878.82^†^ | 582 |
| Middle Cingulate Cortex | M | 2 | 4 | 32 | 572.57^‡^ | 18 |
| Ventromedial Prefrontal Cortex [VMPFC] | M | 0 | 30 | -26 | 546.25^‡^ | 7 |
| ***Cohort 1, [(PF > cPF) > (PL > cPL)]_Contr_* > *[(PF > cPF) > (PL > cPL)]_Med4_*** | | | | | | |
| Anterior Insula [AI] | R | 30 | 16 | -18 | 149.49^§^ | 17 |
| ***Cohort 1, [(PF > cPF) > (PL > cPL)]_Med1_* > *[(PF > cPF) > (PL > cPL)]_Med4_*** | | | | | | |
| Thalamus | R | 8 | -2 | 0 | 851.94^‡^ | 33 |
| ***Cohort 2, Main Effect: (PF > cPF) > (PL > cPL)*** | | | | | | |
| Posterior Insula [PI] | R | 42 | -2 | 4 | 1046.45^‡^ | 52 |
| Amygdala | R | 24 | -4 | -18 | 1722.79^†^ | 250 |
| Amygdala | L | -24 | -2 | -20 | 1367.57^‡^ | 126 |
| Inferior Frontal Gyrus [IFG] | R | 48 | 40 | 10 | 930.19^‡^ | 13 |
| Orbitofrontal Cortex | R | 28 | 32 | -18 | 967.11^‡^ | 23 |
| Precentral Gyrus | L | -46 | 2 | 30 | 1123.61^‡^ | 160 |
| Supramarginal/Postcentral Gyrus [SMG/PCG] | R | 62 | -22 | 32 | 2751.67^†^ | 703 |
| Supramarginal/Postcentral Gyrus [SMG/PCG] | L | -64 | -24 | 28 | 1077.26^‡^ | 39 |
| Intraparietal Sulcus | R | 24 | -64 | 46 | 909.28^‡^ | 8 |
| Fusiform Gyrus | R | 30 | -46 | -16 | 5563.85^*^ | 7925 |
| Occipital Pole | R | 22 | -84 | -8 | 4575.34^*^ |  |
| Inferior Temporal/Occipital Gyrus | R | 52 | -52 | -12 | 2311.52^†^ |  |
| Fusiform Gyrus | L | -28 | -64 | -8 | 5962.31^*^ |  |
| Occipital Pole | L | -12 | -88 | -6 | 7562.57^*^ |  |
| Inferior Temporal/Occipital Gyrus | L | -44 | -56 | -10 | 2816.50^†^ |  |
| ***Cohort 2, [(PF > cPF) > (PL > cPL)] * Post-Graduate Experience*** *(negative effect)* | | | | | | |
| Anterior Insula [AI] | L | -42 | 16 | -10 | 127.27^§^ | 14 |
| * *p* < 0.001; † *p* < 0.01; ‡ *p* < 0.05 FWE corrected for multiple comparisons for the whole brain.  § *p* < 0.05 Small Volume Corrected for multiple comparisons | | | | | | |

##### Supplementary Table S5

Vicarious Pain Signature for Handedness Task. Results from Linear Mixed Model analysis with EMOTIONAL AROUSAL (*Neutral, Negative*) and STIMULI (*Painful, Painless*) as within-subjects factor. In Cohort 1, GROUP (*Controls Med1, Med4*) was modeled as between-subjects factor. In Cohort 2, postgraduate EXPERIENCE was modeled as continuous predictor. The *lmer-syntax* of the tested model is the following:

Cohort 1 – *Dep. Var. ~ EA*STIMULI*GROUP + (EA+STIMULI|Subjects)*.

Cohort 2 – *Dep. Var. ~ EA*STIMULI*EXP + (EA+STIMULI|Subjects)*.

Differently from behavioral measures, the analysis was not carried out on single-trial data, but on parameter-maps from first-level neuroimaging analysis, obtained by collapsing all trials of each condition/subject together. As such, we could not account for random effects of Materials. Furthermore, the model could be fit under a simplified random structure for the effect of subjects’ identity. The analysis was run on the output of two Vicarious Pain models: *Krishan_2016_* and *Zhou-NS_2020_*. For each of these, and for each effect of interest, the table reports the *t*-value associated with the model parameters. Significant effects are highlighted.

|  | ***Krishnan_2016_*** | ***Zhou-NS_2020_*** |
| --- | --- | --- |
| ***COHORT 1*** |  |  |
| *EMOTIONAL AROUSAL* [EA] | *t*_(79)_ = **-2.67**^‡^ | *t*_(118)_ = **-2.55**^‡^ |
| *STIMULI* | *t*_(76)_ = -0.26 | *t*_(118)_ = 0.15 |
| *GROUP* [GR]*: Contr. vs. Med1* | *t*_(42)_ = 0.24 | *t*_(42)_ = 0.28 |
| *GROUP* [GR]*: Contr. vs. Med4* | *t*_(42)_ = -0.30 | *t*_(42)_ = -0.70 |
| *EA*STIMULI* | *t*_(140)_ = 1.21 | *t*_(120)_ = 0.66 |
| *EA*GR: Contr. vs. Med1* | *t*_(79)_ = 0.11 | *t*_(119)_ = 1.23 |
| *EA*GR: Contr. vs. Med4* | *t*_(79)_ = 0.06 | *t*_(118)_ = **-2.37**^‡^ |
| *STIMULI*GR: Contr. vs. Med1* | *t*_(76)_ = -0.98 | *t*_(118)_ = **-2.23**^‡^ |
| *STIMULI*GR: Contr. vs. Med4* | *t*_(76)_ = -0.95 | *t*_(118)_ = **-2.57**^‡^ |
| *EA*STIMULI*GR: Contr. vs. Med1* | *t*_(40)_ = 0.51 | *t*_(120)_ = -1.18 |
| *EA*STIMULI*GR: Contr. vs. Med4* | *t*_(40)_ = 0.32 | *t*_(118)_ = **-2.85**^†^ |
| ***COHORT 2*** |  |  |
| *EMOTIONAL AROUSAL* [EA] | *t*_(64)_ = -**6.44^*^** | *t*_(84)_ = 1.44 |
| *STIMULI* | *t*_(63)_ = -**4.59^*^** | *t*_(84)_ = **4.50^*^** |
| *EXPERIENCE* [EXP] | *t*_(29)_ = 0.81 | *t*_(30)_ = 0.23 |
| *EA*STIMULI* | *t*_(56)_ = **6.03^*^** | *t*_(84)_ = -**3.53^*^** |
| *EA*EXP* | *t*_(84)_ = 0.71 | *t*_(84)_ = 0.31 |
| *STIMULI*EXP* | *t*_(79)_ = -1.51 | *t*_(79)_ = 0.71 |
| *EA*STIMULI*EXP* | *t*_(84)_ = 0.06 | *t*_(84)_ = -1.76 |
| **p < 0.001; †p < 0.01; ‡p < 0.05* | | |

##### Supplementary Table S6

Cognitive and Affective Theory of Mind Task. Results from (Generalized) Linear Mixed Model analysis with STORY CATEGORY (*B, E, Pa, Ph*) as within-subjects factor. In Cohort 1, GROUP (*Controls Med1, Med4*) was modeled as between-subjects factor. In Cohort 2, postgraduate EXPERIENCE was modeled as continuous predictor. The *lmer-syntax* of the tested model is the following:

Cohort 1 *– Dep. Var. ~ STORY*GROUP + (STORY|Subjects) + (GROUP|Materials)*.

Cohort 2 – *Dep. Var. ~ STORY*EXP + (STORY|Subjects) + (EXP|Materials)*.

The analysis was run on Accuracy and Reaction Times of correct responses. For each dependent variable (displayed vertically), and for each effect of interest (horizontally), the table reports the *Z*/*t*-value associated with the model parameters.

|  | ***Accuracy*** | ***RTs*** *[sec]* |
| --- | --- | --- |
| ***COHORT 1*** |  |  |
| *STORY CATEGORY* [SC]: *Photos* [Ph] vs. *Beliefs* [B] | *Z* = 1.21 | *t*_(46)_ = -0.43 |
| *SC: Ph* vs. *Emotions* [E] | *Z* = 0.79 | *t*_(51)_ = 0.61 |
| *SC: Ph* vs.*Pain* [Pa] | *Z* = 0.88 | *t*_(47)_ = 1.64 |
| *GROUP* [GR]*: Contr. vs. Med1* | *Z* = -0.34 | *t*_(43)_ = 1.18 |
| *GROUP* [GR]*: Contr. vs. Med4* | *Z* = 0.62 | *t*_(43)_ = 1.03 |
| *SC*GR*: (*Ph* vs. *B*)*(*Contr. vs. Med1*) | *Z* = 1.11 | *t*_(41)_ = -0.69 |
| *SC*GR*: (*Ph* vs. *E*)*(*Contr. vs. Med1*) | *Z* = -0.88 | *t*_(39)_ = 0.56 |
| *SC*GR*: (*Ph* vs. *Pa*)*(*Contr. vs. Med1*) | *Z* = 0.61 | *t*_(40)_ = 0.29 |
| *SC*GR*: (*Ph* vs. *B*)*(*Contr. vs. Med4*) | *Z* = 0.30 | *t*_(40)_ = -0.96 |
| *SC*GR*: (*Ph* vs. *E*)*(*Contr. vs. Med4*) | *Z* = -0.31 | *t*_(39)_ = -0.05 |
| *SC*GR*: (*Ph* vs. *Pa*)*(*Contr. vs. Med4*) | *Z* = 0.63 | *t*_(39)_ = -0.35 |
| ***COHORT 2*** |  |  |
| *STORY CATEGORY* [SC]: *Photos* [Ph] vs. *Beliefs* [B] | *Z* = 1.28 | *t*_(44)_ = -0.65 |
| *SC: Ph* vs. *Emotions* [E] | *Z* = 0.05 | *t*_(43)_ = 0.66 |
| *SC: Ph* vs.*Pain* [Pa] | *Z* = 1.12 | *t*_(48)_ = 1.88 |
| *EXPERIENCE* [EXP] | *Z* = 1.18 | *t*_(26)_ = 0.50 |
| *SC*EXP*: (*Ph* vs. *B*) | *Z* = -0.12 | *t*_(37)_ = 0.72 |
| *SC*EXP*: (*Ph* vs. *E*) | *Z* = -0.46 | *t*_(89)_ = 1.37 |
| *SC*EXP*: (*Ph* vs. *Pa*) | *Z* = -1.07 | *t*_(27)_ = 0.55 |
| **p < 0.001; †p < 0.01; ‡p < 0.05* | | |

##### Supplementary Table S7

Cognitive and Affective Theory of Mind Task. Regions displaying suprathreshold activity evoked by the attribution of *beliefs* while reading the text-based *Scenarios*.

|  | ***SIDE*** | ***Coordinates*** | | | ***TFCE*** | ***Cluster size*** |
| --- | --- | --- | --- | --- | --- | --- |
|  |  | ***x*** | ***y*** | ***z*** |  |  |
| ***Cohort 1, Main Effect: Attribution of Beliefs*** | | | | | | |
| Temporo-Parietal Junction [TPJ] | R | 58 | -54 | 28 | 9136.88^*^ | 18590 |
| Middle Temporal Gyrus | R | 58 | -28 | -8 | 3028.99^†^ |  |
| Dorsolateral Prefrontal Cortex [DLPFC] | R | 40 | 20 | 36 | 2591.07^†^ |  |
| Dorsolateral Prefrontal Cortex [DLPFC] | L | -40 | 18 | 40 | 1967.04^†^ |  |
| Inferior Frontal Gyrus | R | 50 | 28 | -10 | 1576.62^‡^ |  |
| Medial Prefrontal Cortex *(dorsal part)* | M | 4 | 42 | 44 | 5157.80^*^ |  |
| Medial Prefrontal Cortex *(rostral part)* | M | 2 | 60 | 12 | 6921.89^*^ |  |
| Temporo-Parietal Junction [TPJ] | L | -54 | -60 | 26 | 6317.21^*^ | 1980 |
| Middle Temporal Gyrus | L | -62 | -22 | -14 | 3936.84^†^ | 569 |
| Temporal Pole | L | -46 | 8 | -36 | 450.50^‡^ | 109 |
| Inferior Frontal Gyrus | L | -40 | 22 | -22 | 613.81^‡^ | 344 |
| Temporal Pole | L | -34 | 16 | -28 | 451.29^‡^ |  |
| Caudate | R | 14 | 2 | 12 | 438.55^‡^ | 6898 |
| Caudate | L | -10 | 10 | 2 | 553.20^‡^ |  |
| Precuneus | M | 2 | -60 | 38 | 17576.89^*^ |  |
| Posterior Cingulate Cortex | M | -6 | -46 | 6 | 840.03^†^ |  |
| Thalamus | M | 0 | -4 | 4 | 463.94^‡^ |  |
| ***Cohort 2, Main Effect: Attribution of Beliefs*** | | | | | | |
| Temporo-Parietal Junction [TPJ] | R | 50 | -52 | 22 | 1280.61^‡^ | 225 |
| Precuneus | M | -4 | -58 | 38 | 1775.65^†^ | 663 |
| * *p* < 0.001; † *p* < 0.01; ‡ *p* < 0.05 corrected for multiple comparisons at the cluster level for the whole. | | | | | | |

##### Supplementary Table S8

Cognitive and Affective Theory of Mind Task. Regions displaying suprathreshold activity evoked by the attribution of *emotions* while reading the text-based *Scenarios*.

|  | ***SIDE*** | ***Coordinates*** | | | ***TFCE*** | ***Cluster size*** |
| --- | --- | --- | --- | --- | --- | --- |
|  |  | ***x*** | ***y*** | ***z*** |  |  |
| ***Cohort 1, Main Effect: Attribution of Emotions*** | | | | | | |
| Middle-Posterior Insula | R | 40 | 2 | 10 | 1927.11^*^ | 48875 |
| Middle-Posterior Insula | L | -34 | 2 | 8 | 2640.99^*^ |  |
| Medial Prefrontal Cortex *(ventral part)* | M | -4 | 34 | -8 | 2068.62^*^ |  |
| Superior Temporal Gyrus | R | 66 | -14 | 0 | 1252.68^†^ |  |
| Superior Temporal Gyrus | L | -56 | -4 | 14 | 3145.19^*^ |  |
| Middle Cingulate Cortex | M | -2 | -16 | 40 | 1932.39^*^ |  |
| Precentral Gyrus | R | 50 | -4 | 52 | 1387.84^†^ |  |
| Postcentral Gyrus | R | 38 | -30 | 58 | 2759.07^*^ |  |
| Postcentral Gyrus | L | -36 | -28 | 56 | 2568.22^*^ |  |
| Supramarginal Gyrus | R | 62 | -30 | 20 | 3053.89^*^ |  |
| Supramarginal Gyrus | L | -58 | -28 | 22 | 2804.15^*^ |  |
| Intraparietal Sulcus | R | 20 | -48 | 64 | 3113.26^*^ |  |
| Intraparietal Sulcus | L | -18 | -50 | 62 | 2385.48^*^ |  |
| Inferior Temporal/Occipital Gyrus | R | 50 | -70 | -2 | 2915.07^*^ |  |
| Inferior Temporal/Occipital Gyrus | L | -48 | -68 | 4 | 2369.84^*^ |  |
| Fusiform Gyrus | R | 38 | -54 | -16 | 2979.78^*^ |  |
| Occipital Pole | R | 22 | -90 | 16 | 3548.23^*^ |  |
| Occipital Pole | L | -38 | -86 | 0 | 3338.35^*^ |  |
| Cerebellum | R | 30 | -58 | -24 | 3408.53^*^ |  |
| Cerebellum | L | -8 | -60 | -14 | 1995.78^*^ |  |
| Temporal Pole | R | 28 | 10 | -22 | 983.81^‡^ | 141 |
| Precentral Gyrus | L | -52 | -4 | 44 | 886.74^‡^ | 123 |
| ***Cohort 2, Main Effect: Attribution of Emotions*** | | | | | | |
| Posterior Insula [PI] | R | 40 | -4 | 4 | 976.60^‡^ | 257 |
| Superior Temporal Gyrus | R | 64 | -34 | 18 | 1178.21^‡^ | 1050 |
| Supramarginal Gyrus | R | 62 | -20 | 42 | 1240.44^‡^ |  |
| Anterior Insula [AI] | L | -36 | 6 | 4 | 1214.95^‡^ | 2175 |
| Superior Temporal Gyrus | L | -56 | -8 | -4 | 1542.96^‡^ |  |
| Parietal Operculum | L | -52 | -38 | 24 | 1091.44^‡^ |  |
| Medial Prefrontal Cortex *(ventral part)* | M | 4 | 40 | 6 | 1391.21^‡^ | 684 |
| Precuneus | M | 10 | -44 | 62 | 1054.36^‡^ | 972 |
| Precentral Gyrus | R | 28 | -16 | 74 | 1250.19^‡^ |  |
| Postcentral Gyrus | R | 46 | -16 | 62 | 963.97^‡^ | 1 |
| Precentral Gyrus | L | -18 | -20 | 76 | 1313.71^‡^ | 3287 |
| Postcentral Gyrus | L | -18 | -46 | 72 | 1523.69^†^ |  |
| Middle Cingulate Cortex | M | -4 | -18 | 40 | 1308.80^‡^ |  |
| * *p* < 0.001; † *p* < 0.01; ‡ *p* < 0.05 corrected for multiple comparisons at the cluster level for the whole. | | | | | | |

##### Supplementary Table S9

Cognitive and Affective Theory of Mind Task. Regions displaying suprathreshold activity evoked by the attribution of *pain* while reading the text-based *Scenarios*.

|  | ***SIDE*** | ***Coordinates*** | | | ***TFCE*** | ***Cluster size*** |
| --- | --- | --- | --- | --- | --- | --- |
|  |  | ***x*** | ***y*** | ***z*** |  |  |
| ***Cohort 1, Main Effect: Attribution of Pain*** | | | | | | |
| Posterior Insula [PI] | R | 40 | 2 | -10 | 883.03^‡^ | 218 |
| Anterior Insula [AI] | R | 48 | 14 | -2 | 651.75^‡^ |  |
| Posterior Insula [PI] | L | -38 | -14 | -6 | 1143.43^‡^ | 369 |
| Anterior Insula [AI] | L | -36 | 12 | -2 | 955.16^‡^ |  |
| Supramarginal/Postcentral Gyrus | L | -56 | -36 | 52 | 3591.43^*^ | 1536 |
| Supramarginal/Postcentral Gyrus | R | 60 | -36 | 36 | 2631.26^†^ | 806 |
| Inferior Frontal Gyrus | R | 44 | 42 | 2 | 2341.26^†^ | 416 |
| Dorsolateral Prefrontal Cortex | L | -26 | 34 | 46 | 1445.24^†^ | 18229 |
| Inferior Frontal Gyrus | L | -46 | 40 | 8 | 3649.57^*^ |  |
| Posterior Orbital Gyrus | L | -24 | 30 | -14 | 6272.60^*^ |  |
| Precuneus | M | -2 | -74 | 46 | 5097.21^*^ |  |
| Posterior Cingulate Cortex | M | -8 | -30 | 40 | 8757.16^*^ |  |
| Middle Cingulate Cortex [MCC] | M | -4 | 2 | 36 | 5374.55^*^ |  |
| Medial Prefrontal Cortex (*ventral part*) | M | 4 | 50 | -6 | 1326.54^†^ |  |
| ***Cohort 1, Pain_Contr_ > Pain_Med4_*** | | | | | | |
| Anterior Insula [AI] | R | 42 | 16 | -4 | 205.90^§^ | 131 |
| * *p* < 0.001; † *p* < 0.01; ‡ *p* < 0.05 corrected for multiple comparisons at the cluster level for the whole.  § *p* < 0.05 Small Volume Corrected for multiple comparisons. | | | | | | |

##### Supplementary Table S10

Cognitive and Affective Theory of Mind Task. Regions displaying suprathreshold activity evoked by making active *judgment* about *beliefs & emotions*, compared to the control *photos* condition.

|  | ***SIDE*** | ***Coordinates*** | | | ***TFCE*** | ***Cluster size*** |
| --- | --- | --- | --- | --- | --- | --- |
|  |  | ***x*** | ***y*** | ***z*** |  |  |
| ***Cohort 1, Main Effect: Beliefs > Photos Judgment*** | | | | | | |
| Temporo-Parietal Junction [TPJ] | R | 52 | -48 | 16 | 956.38^‡^ | 182 |
| Middle Temporal Gyrus | R | 60 | -24 | -10 | 1318.29^‡^ | 117 |
| Precuneus | M | 6 | -60 | 34 | 6314.94^*^ | 4520 |
| Posterior Cingulate Cortex | M | -2 | -20 | 32 | 929.46^‡^ |  |
| ***Cohort 2, Main Effect: Beliefs > Photos Judgment*** | | | | | | |
| Precuneus | M | 0 | -52 | 32 | 1577.45^‡^ | 491 |
| ***Cohort 1, Main Effect: Emotions > Photos Judgment*** | | | | | | |
| Precuneus | M | 2 | -68 | 36 | 1259.75^†^ | 938 |
| Posterior Cingulate Cortex | M | -2 | -48 | 30 | 2608.71^*^ |  |
| Middle Cingulate Cortex | M | 2 | -14 | 40 | 1837.41^†^ | 477 |
| Medial Prefrontal Cortex *(dorsal part)* | M | -10 | 62 | 26 | 1904.68^†^ | 570 |
| Fusiform Gyrus | L | -34 | -54 | -20 | 1172.38^‡^ | 79 |
| Inferior Frontal Gyrus | L | -42 | 22 | -20 | 1114.92^‡^ | 136 |
| Amygdala | L | -20 | 0 | -14 | 890.48^‡^ | 29 |
| Putamen | L | -16 | 14 | 0 | 964.23^‡^ | 103 |
| ***Cohort 1, (Emotions > Photos)_Med1_ > (Emotions > Photos)_Med4_ Judgment*** | | | | | | |
| dorsal Anterior Cingulate Cortex [dACC] | M | 0 | 36 | 28 | 97.87^§^ | 8 |
| ***Cohort 1, Main Effect: Pain > Photos Judgment*** | | | | | | |
| Posterior Insula [PI] | R | 40 | -8 | -10 | 1494.14^†^ | 14426 |
| Posterior Insula [PI] | L | -36 | -8 | -10 | 1485.93^†^ |  |
| Anterior Insula [AI] | R | 42 | 4 | -4 | 1598.48^†^ |  |
| Anterior Insula [AI] | L | -36 | 8 | -4 | 1332.05^†^ |  |
| Putamen | R | 20 | 12 | -6 | 1879.08^*^ |  |
| Putamen | L | -18 | 12 | -10 | 2287.48^*^ |  |
| Precentral Gyrus | L | -38 | -14 | 56 | 1486.51^†^ |  |
| Supramarginal/Postcentral Gyrus | R | 66 | -34 | 20 | 1168.58^†^ |  |
| Supramarginal/Postcentral Gyrus | L | -64 | -30 | 22 | 1458.14^†^ |  |
| Middle Cingulate Cortex | M | 6 | -20 | 34 | 1469.19^†^ |  |
| Supplementary Motor Area | M | 10 | 6 | 62 | 1185.37^†^ |  |
| Precentral Gyrus | R | 46 | -4 | 48 | 786.33^‡^ | 10 |
| Lingual Gyrus | R | 18 | -70 | 0 | 2183.70^*^ | 12188 |
| Lingual Gyrus | L | 10 | -60 | 2 | 1047.87^†^ |  |
| Fusiform Gyrus | R | 34 | -58 | -20 | 2068.63^*^ |  |
| Fusiform Gyrus | L | -34 | -58 | -18 | 1327.94^†^ |  |
| Inferior Occipital Gyrus | R | 40 | -76 | 6 | 1983.60^*^ |  |
| Inferior Occipital Gyrus | L | -46 | -76 | -2 | 1553.54^†^ |  |
| Superior Occipital Gyrus | R | 28 | -80 | 32 | 2016.36^*^ |  |
| Superior Occipital Gyrus | L | -28 | -82 | 24 | 1284.39^†^ |  |
| Superior Parietal Cortex | R | 16 | -72 | 48 | 1963.29^*^ |  |
| Superior Parietal Cortex | L | -12 | -60 | 58 | 1019.07^†^ |  |
| Thalamus | R | 10 | -24 | -2 | 1051.05^†^ |  |
| Thalamus | L | -10 | -24 | 0 | 997.60^‡^ |  |
| Precuneus | M | 16 | -72 | 36 | 1926.88^*^ |  |
| * *p* < 0.001; † *p* < 0.01; ‡ *p* < 0.05 corrected for multiple comparisons at the cluster level for the whole.  § *p* < 0.05 Small Volume Corrected for multiple comparisons. | | | | | | |

##### Supplementary Table S11

Representational Similarity Analysis. Regions encoding pain-specific information in the two cohorts (within-pain *vs*. across-affect).

|  | ***SIDE*** | ***Coordinates*** | | | ***TFCE*** | ***Cluster size*** |
| --- | --- | --- | --- | --- | --- | --- |
|  |  | ***x*** | ***y*** | ***z*** |  |  |
| ***Cohort 1, Main effect*** | | | | | | |
| Inferior Frontal Gyrus [IFG] | R | 48 | 36 | 0 | 1761.90^†^ | 790 |
| Anterior Insula [AI] | L | -32 | 24 | 8 | 1384.81^‡^ | 2325 |
| Inferior Frontal Gyrus [IFG] | L | -42 | 36 | 10 | 2055.33^†^ |  |
| Orbitofrontal Cortex | L | -22 | 34 | -10 | 2426.10^*^ |  |
| Middle Insula [MI] | M | -40 | 6 | 2 | 1197.04^‡^ | 619 |
| ***Cohort 2, Main effect*** | | | | | | |
| Hippocampus [Hipp] | R | 32 | -10 | -20 | 1217.10^‡^ | 727 |
| Amygdala [Amy] | R | 24 | -6 | -22 | 1217.10^‡^ |  |
| Putamen | R | 28 | -10 | -8 | 1248.20^‡^ |  |
| Temporal Pole | R | 48 | 0 | -38 | 1317.69^‡^ |  |
| Hippocampus [Hipp] | L | -20 | -14 | -22 | 1263.41^‡^ | 357 |
| Amygdala [Amy] | L | -18 | -8 | -16 | 1229.51^‡^ |  |
| Superior Frontal Gyrus | R | 24 | 12 | 60 | 1230.61^‡^ | 282 |
| dorsal Anterior Cingulate Cortex [dACC] | M | 0 | 34 | 14 | 117.03^§^ | 20 |
| Supplementary Motor Area | M | 4 | 24 | 54 | 1201.56^‡^ | 109 |
| Cerebellum | R | 26 | -76 | -38 | 1351.70^‡^ | 317 |
| * *p* < 0.001; † *p* < 0.01; ‡ *p* < 0.05 FWE corrected for multiple comparisons for the whole brain.  § *p* < 0.05 Small Volume Corrected for multiple comparisons. | | | | | | |

##### Supplementary Table S12

Quantitative comparison between Cohorts 1 & 2: Behavioral results from the Handedness Task. Results from (Generalized) Linear Mixed Model analysis with EMOTIONAL AROUSAL (*Neutral, Negative*) and STIMULI (*Painful, Painless*) as within-subjects factor, and GROUP (*Controls Med1, Med4, Nurses*) as between-subjects factor. The *lmer-syntax* of the tested model is the following:

*Dep. Var. ~ EA*STIMULI*GROUP + EA*STIMULI*AGE + (EA*STIMULI|Subjects) + (GROUP + AGE|Materials)*.

As cohort 1 included only females, this analysis was carried out by including only the females from cohort 2. Furthermore, as cohort 2 underwent overall shorter version of the paradigm we considered only a sub-selection of the data (120 trials out of 180) from cohort 1 in order to match the two datasets. For each dependent variable (displayed vertically), and for each effect of interest (horizontally), the table reports the *Z*/*t*-value associated with the model parameters. Significant effects are highlighted

|  | ***Accuracy*** | ***RTs*** *[sec]* | ***Arousal*** *[1; 10]* | ***Valence*** *[-4; 4]* | ***Pain*** *[1; 10]* | ***Familiarity*** *[1; 10]* |
| --- | --- | --- | --- | --- | --- | --- |
| *EMOTIONAL AROUSAL* [EA] | *Z* = **-3.65^*^** | *t*_(119)_ = **2.57^‡^** | *t*_(82)_ = **9.87^*^** | *t*_(107)_ = -**7.91^*^** | *t*_(75)_ = -**13.52^*^** | *t*_(102)_ = -**3.97^*^** |
| *STIMULI* | *Z* = -1.25 | *t*_(115)_ = 1.25 | *t*_(111)_ = 1.74 | *t*_(136)_ = **2.51^‡^** | *t*_(129)_ = -0.28 | *t*_(105)_ = 1.34 |
| *GROUP* [GR]*: Contr. vs. Med1* | *Z* = 0.02 | *t*_(62)_ = 0.86 | *t*_(58)_ = 1.24 | *t*_(58)_ = 1.26 | *t*_(58)_ = 0.48 | *t*_(60)_ = 1.11 |
| *GROUP* [GR]*: Contr. vs. Med4* | *Z* = 0.56 | *t*_(58)_ = 0.51 | *t*_(60)_ = -0.20 | *t*_(63)_ = 0.99 | *t*_(58)_ = 0.92 | *t*_(61)_ = **4.28^*^** |
| *GROUP* [GR]*: Contr. vs. Nurses* | *Z* = -1.54 | *t*_(59)_ = 0.90 | *t*_(60)_ = 0.04 | *t*_(62)_ = 1.22 | *t*_(61)_ = **2.64^‡^** | *t*_(66)_ = **5.02^*^** |
| *AGE* | *Z* = 1.26 | *t*_(61)_ = 0.38 | *t*_(59)_ = -0.52 | *t*_(59)_ = -0.21 | *t*_(59)_ = -0.65 | *t*_(60)_ = 0.87 |
| *EA*STIMULI* | *Z* = 0.31 | *t*_(122)_ = -0.33 | *t*_(88)_ = **-4.82^*^** | *t*_(129)_ = 0.65 | *t*_(72)_ = **-8.04^*^** | *t*_(122)_ = 0.20 |
| *EA*GR: Contr. vs. Med1* | *Z* = 1.22 | *t*_(72)_ = -1.41 | *t*_(59)_ = **-2.49^‡^** | *t*_(59)_ = -0.01 | *t*_(59)_ = -0.84 | *t*_(60)_ = 0.66 |
| *EA*GR: Contr. vs. Med4* | *Z* = 0.55 | *t*_(61)_ = -1.93 | *t*_(61)_ = -1.92 | *t*_(64)_ = 1.06 | *t*_(58)_ = -0.62 | *t*_(63)_ = -0.88 |
| *EA*GR: Contr. vs. Nurses* | *Z* = 1.60 | *t*_(64)_ = -1.49 | *t*_(61)_ = -0.01 | *t*_(62)_ = 1.17 | *t*_(62)_ = -1.10 | *t*_(73)_ = -1.04 |
| *STIMULI*GR: Contr. vs. Med1* | *Z* = 0.33 | *t*_(104)_ = 0.67 | *t*_(59)_ = 0.07 | *t*_(60)_ = -1.50 | *t*_(59)_ = -0.66 | *t*_(60)_ = -0.23 |
| *STIMULI*GR: Contr. vs. Med4* | *Z* = 0.27 | *t*_(279)_ = -1.85 | *t*_(64)_ = 1.35 | *t*_(70)_ = **-2.24^‡^** | *t*_(59)_ = -1.17 | *t*_(63)_ = **-4.57^*^** |
| *STIMULI*GR: Contr. vs. Nurses* | *Z* = 0.94 | *t*_(213)_ = **-2.34^‡^** | *t*_(64)_ = 1.51 | *t*_(67)_ = **-2.02^‡^** | *t*_(74)_ = -1.46 | *t*_(73)_ = **-4.64^*^** |
| *EA*AGE* | *Z* = -0.87 | *t*_(70)_ = 1.27 | *t*_(60)_ = **-2.21^‡^** | *t*_(59)_ = 0.49 | *t*_(59)_ = -0.27 | *t*_(61)_ = 1.02 |
| *STIMULI*AGE* | *Z* = -0.92 | *t*_(109)_ = **2.09^‡^** | *t*_(61)_ = -0.13 | *t*_(60)_ = -0.75 | *t*_(60)_ = 0.41 | *t*_(61)_ = -0.55 |
| *EA*STIM*GR: Contr. vs. Med1* | *Z* = -0.48 | *t*_(77)_ = 1.46 | *t*_(60)_ = 1.67 | *t*_(59)_ = -0.33 | *t*_(58)_ = 0.57 | *t*_(61)_ = -0.86 |
| *EA*STIM*GR: Contr. vs. Med4* | *Z* = -0.46 | *t*_(66)_ = **2.17^‡^** | *t*_(62)_ = 1.62 | *t*_(68)_ = -0.67 | *t*_(58)_ = 0.15 | *t*_(64)_ = 0.91 |
| *EA*STIM*GR: Contr. vs. Nurses* | *Z* = -0.36 | *t*_(70)_ = 0.30 | *t*_(62)_ = 1.92 | *t*_(65)_ = **-2.12^‡^** | *t*_(61)_ = 1.47 | *t*_(80)_ = 1.18 |
| *EA*STIM*AGE* | *Z* = 0.37 | *t*_(75)_ = -0.38 | *t*_(60)_ = 1.18 | *t*_(59)_ = 0.13 | *t*_(59)_ = -0.06 | *t*_(62)_ = -1.47 |
| **p < 0.001; †p < 0.01; ‡p < 0.05* | | | | | | |

##### Supplementary Table S13

Quantitative comparison between Cohorts 1 & 2: Cognitive and Affective Theory of Mind Task. Results from (Generalized) Linear Mixed Model analysis with STORY CATEGORY (*B, E, Pa, Ph*) as within-subjects factor, and GROUP (*Controls Med1, Med4, Nurses*) as between-subjects factor. The *lmer-syntax* of the tested model is the following:

*Dep. Var. ~ STORY*GROUP + STORY*AGE + (STORY|Subjects) + (GROUP + AGE|Materials)*.

As cohort 1 included only females, this analysis was carried out by including only the females from cohort 2. As cohort 2 underwent only one session, we included only the first session (out of two) from cohort 1 in order to match the two datasets. The analysis was run on Accuracy and Reaction Times of correct responses. For each dependent variable (displayed vertically), and for each effect of interest (horizontally), the table reports the *Z*/*t*-value associated with the model parameters.

|  | ***Accuracy*** | ***RTs*** *[sec]* |
| --- | --- | --- |
| *STORY CATEGORY* [SC]: *Photos* [Ph] vs. *Beliefs* [B] | *Z* = 0.53 | *t*_(44)_ = -0.20 |
| *SC: Ph* vs. *Emotions* [E] | *Z* = 0.47 | *t*_(44)_ = 0.35 |
| *SC: Ph* vs.*Pain* [Pa] | *Z* = 0.35 | *t*_(43)_ = 1.49 |
| *GROUP* [GR]*: Contr. vs. Med1* | *Z* = 0.56 | *t*_(56)_ = 1.06 |
| *GROUP* [GR]*: Contr. vs. Med4* | *Z* = 0.74 | *t*_(58)_ = 0.94 |
| *GROUP* [GR]*: Contr. vs. Nurses* | *Z* = -1.04 | *t*_(55)_ = 0.44 |
| *AGE* | *Z* = 0.86 | *t*_(55)_ = 0.54 |
| *SC*GR*: (*Ph* vs. *B*)*(*Contr. vs. Med1*) | *Z* = -0.11 | *t*_(45)_ = -0.34 |
| *SC*GR*: (*Ph* vs. *E*)*(*Contr. vs. Med1*) | *Z* = -1.17 | *t*_(43)_ = 1.80 |
| *SC*GR*: (*Ph* vs. *Pa*)*(*Contr. vs. Med1*) | *Z* = -0.74 | *t*_(44)_ = 0.32 |
| *SC*GR*: (*Ph* vs. *B*)*(*Contr. vs. Med4*) | *Z* = -0.07 | *t*_(37)_ = -1.26 |
| *SC*GR*: (*Ph* vs. *E*)*(*Contr. vs. Med4*) | *Z* = -0.31 | *t*_(37)_ = -0.19 |
| *SC*GR*: (*Ph* vs. *Pa*)*(*Contr. vs. Med4*) | *Z* = -0.19 | *t*_(35)_ = -0.11 |
| *SC*GR*: (*Ph* vs. *B*)*(*Contr. vs. Nurses*) | *Z* = 0.49 | *t*_(59)_ = -0.04 |
| *SC*GR*: (*Ph* vs. *E*)*(*Contr. vs. Nurses*) | *Z* = 0.28 | *t*_(56)_ = 0.15 |
| *SC*GR*: (*Ph* vs. *Pa*)*(*Contr. vs. Nurses*) | *Z* = 0.61 | *t*_(59)_ = -0.24 |
| *SC*Age*: (*Ph* vs. *B*) | *Z* = -0.60 | *t*_(61)_ = 0.35 |
| *SC*Age*: (*Ph* vs. *E*) | *Z* = -0.84 | *t*_(60)_ = 0.50 |
| *SC*Age*: (*Ph* vs. *Pa*) | *Z* = -0.47 | *t*_(61)_ = 0.58 |
| **p < 0.001; †p < 0.01; ‡p < 0.05* | | |

##### Supplementary Table S14

Quantitative comparison between Cohorts 1 & 2: neuroimaging analysis. Follow-up analysis obtained by combining data from the two cohorts together by assessing effect of healthcare training in terms of four-level GROUP (*Controls*, *Med1*, *Med4*, *Nurses*). As cohort 1 included only females, this analysis was carried out by including only the females from Cohort 2. Furthermore, as cohort 2 underwent overall shorter version of the paradigms we considered only a sub-selection of the data from cohort 1 in order to match the two experiments in terms of power. For the “Handedness” task, we included only those 120 trials (out of 180) which were used in both experiments. For the “Cognitive and Affective Theory of Mind” task, we used data only of the first run in chronological order. Finally, group-level analyses were carried out by including age as nuisance variable of no interest.

|  | ***SIDE*** | ***Coordinates*** | | | ***TFCE*** | ***Cluster size*** |
| --- | --- | --- | --- | --- | --- | --- |
|  |  | ***x*** | ***y*** | ***z*** |  |  |
| ***Handedness Task: (PF > cPF)_Contr_* > *(PF > cPF)_Nurses_*** | | | | | | |
| Anterior Insula [AI] | R | 28 | 10 | -18 | 116.41^§^ | 1 |
| ***Handedness Task: (PF > cPF)_Med1_* > *(PF > cPF)_Nurses_*** | | | | | | |
| Anterior Insula [AI] | R | 32 | 10 | -16 | 62.65^§^ | 5 |
| ***Handedness Task: [(PF > cPF) > (PL > cPL)]_Contr_* > *[(PF > cPF) > (PL > cPL)]_Med4_*** | | | | | | |
| Anterior Insula [AI] | R | 28 | 16 | -14 | 146.93^§^ | 13 |
| ***Cognitive and Affective Theory of Mind Task: Attribution of Pain_Contr_ > Pain_Nurses_*** | | | | | | |
| Anterior Insula | R | 44 | 16 | -2 | 214.68^§^ | 104 |
| § *p* < 0.05 Small Volume Corrected for multiple comparisons | | | | | | |

##### Supplementary Table S15

Quantitative comparison between Cohorts 1 & 2: Vicarious Pain Signature for Handedness Task. Results from Linear Mixed Model analysis with EMOTIONAL AROUSAL (*Neutral, Negative*) and STIMULI (*Painful, Painless*) as within-subjects factor, and GROUP (*Controls Med1, Med4, Nurses*) was modeled as between-subjects factor. The *lmer-syntax* of the tested model is the following:

*Dep. Var. ~ EA*STIMULI*GROUP + EA*STIMULI*AGE + (EA+STIMULI|Subjects)*.

As cohort 1 included only females, this analysis was carried out by including only the females from cohort 2. Furthermore, as cohort 2 underwent overall shorter version of the paradigms we considered only a sub-selection of the data from cohort 1 in order to match the two experiments in terms of power.

|  | ***Krishnan_2016_*** | ***Zhou-NS_2020_*** |
| --- | --- | --- |
| *EMOTIONAL AROUSAL* [EA] | *t*_(105)_ = -**4.39**^*^ | *t*_(161)_ = **-3.31**^†^ |
| *STIMULI* | *t*_(105)_ = **-2.53**^‡^ | *t*_(164)_ = -1.25 |
| *GROUP* [GR]*: Contr. vs. Med1* | *t*_(58)_ = 0.35 | *t*_(59)_ = -0.08 |
| *GROUP* [GR]*: Contr. vs. Med4* | *t*_(58)_ = -0.18 | *t*_(59)_ = -1.55 |
| *GROUP* [GR]*: Contr. vs. Nurses* | *t*_(58)_ = -1.73 | *t*_(59)_ = 0.09 |
| *Age* | *t*_(58)_ = 0.50 | *t*_(59)_ = 0.13 |
| *EA*STIMULI* | *t*_(55)_ = **3.34**^†^ | *t*_(165)_ = 1.22 |
| *EA*GR: Contr. vs. Med1* | *t*_(105)_ = 1.29 | *t*_(161)_ = 1.53 |
| *EA*GR: Contr. vs. Med4* | *t*_(105)_ = 0.08 | *t*_(161)_ = **3.68**^*^ |
| *EA*GR: Contr. vs. Nurses* | *t*_(105)_ = 0.12 | *t*_(161)_ = **2.58**^‡^ |
| *STIMULI*GR: Contr. vs. Med1* | *t*_(105)_ = 0.01 | *t*_(164)_ = **2.88**^†^ |
| *STIMULI*GR: Contr. vs. Med4* | *t*_(105)_ = -0.13 | *t*_(164)_ = **3.70**^*^ |
| *STIMULI*GR: Contr. vs. Nurses* | *t*_(105)_ = 0.04 | *t*_(164)_ = 1.44 |
| *EA*AGE* | *t*_(105)_ = 0.84 | *t*_(161)_ = -0.62 |
| *STIMULI*AGE* | *t*_(105)_ = -0.69 | *t*_(164)_ = 0.46 |
| *EA*STIMULI*GR: Contr. vs. Med1* | *t*_(55)_ = -0.43 | *t*_(165)_ = -1.67 |
| *EA*STIMULI*GR: Contr. vs. Med4* | *t*_(55)_ = 0.24 | *t*_(165)_ = **-3.66**^*^ |
| *EA*STIMULI*GR: Contr. vs. Nurses* | *t*_(55)_ = 0.29 | *t*_(165)_ = -1.23 |
| *EA*STIMULI*AGE* | *t*_(55)_ = -0.25 | *t*_(165)_ = -0.59 |
| **p < 0.001; †p < 0.01; ‡p < 0.05* | | |

##### Supplementary Figure S1

Quantitative comparison between Cohorts 1 & 2: Behavioral results from the Handedness Task. Boxplots displaying (A) Response Times of correct responses and (B) post-scanning Valence. Both variables are displayed as differential values between emotional condition and its neutral control. For each boxplot, the horizontal line represents the median value of the distribution, the star represents the average, the box edges refer to the inter-quartile range, and the whiskers represent the data range within 1.5 of the inter-quartile range. Individual data-points are also displayed color coded, with red dots referring to *PF/cPF* stimuli, and blue dots to *NPL/cNPL* stimuli. *Contr.*: Controls; *Med1 & Med4*: university students enrolled at the first/fourth year of medicine. “***” and “*” refer to significant group differences as tested through linear mixed models (see methods) at *p* < 0.001 and *p* < 0.05 respectively.

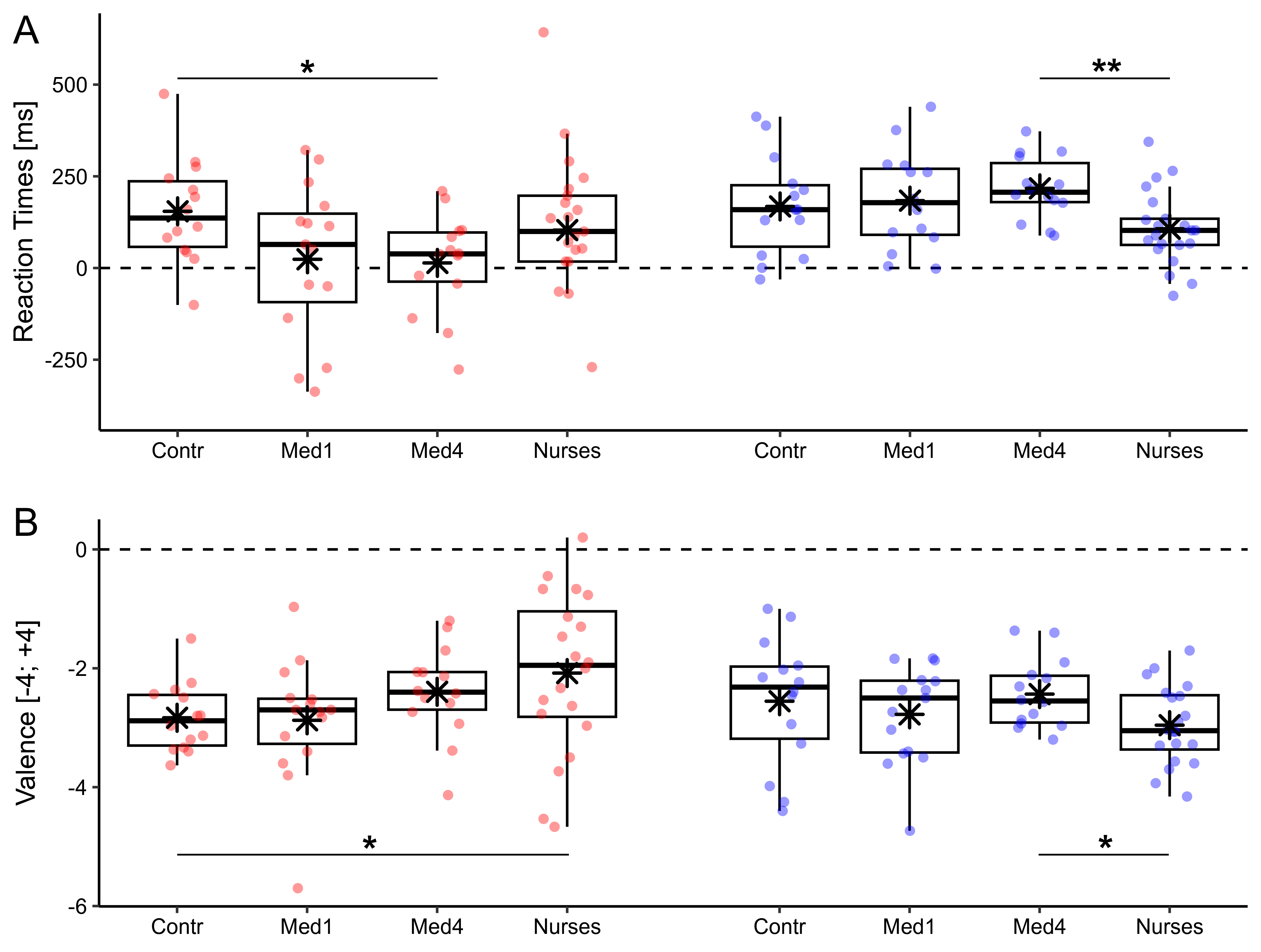

##### Supplementary Figure S2

Adaptation Figure 2 (A), Figure 3 (B) and Figure 4 (B) from the main text to allow for a quantitative comparison between cohorts. As cohort 1 included only females, this analysis was carried out by including only the females from Cohort 2. Furthermore, as cohort 2 underwent overall shorter version of the paradigms we considered only a sub-selection of the data from cohort 1 in order to match the two experiments in terms of power. Renderings are displayed under the same threshold than in the main text. The only exception is subplot A which, for readability purposes, is displayed under *p* < 0.001 (uncorrected), with regions local maxima surviving FWE *p* < 0.05 correction.

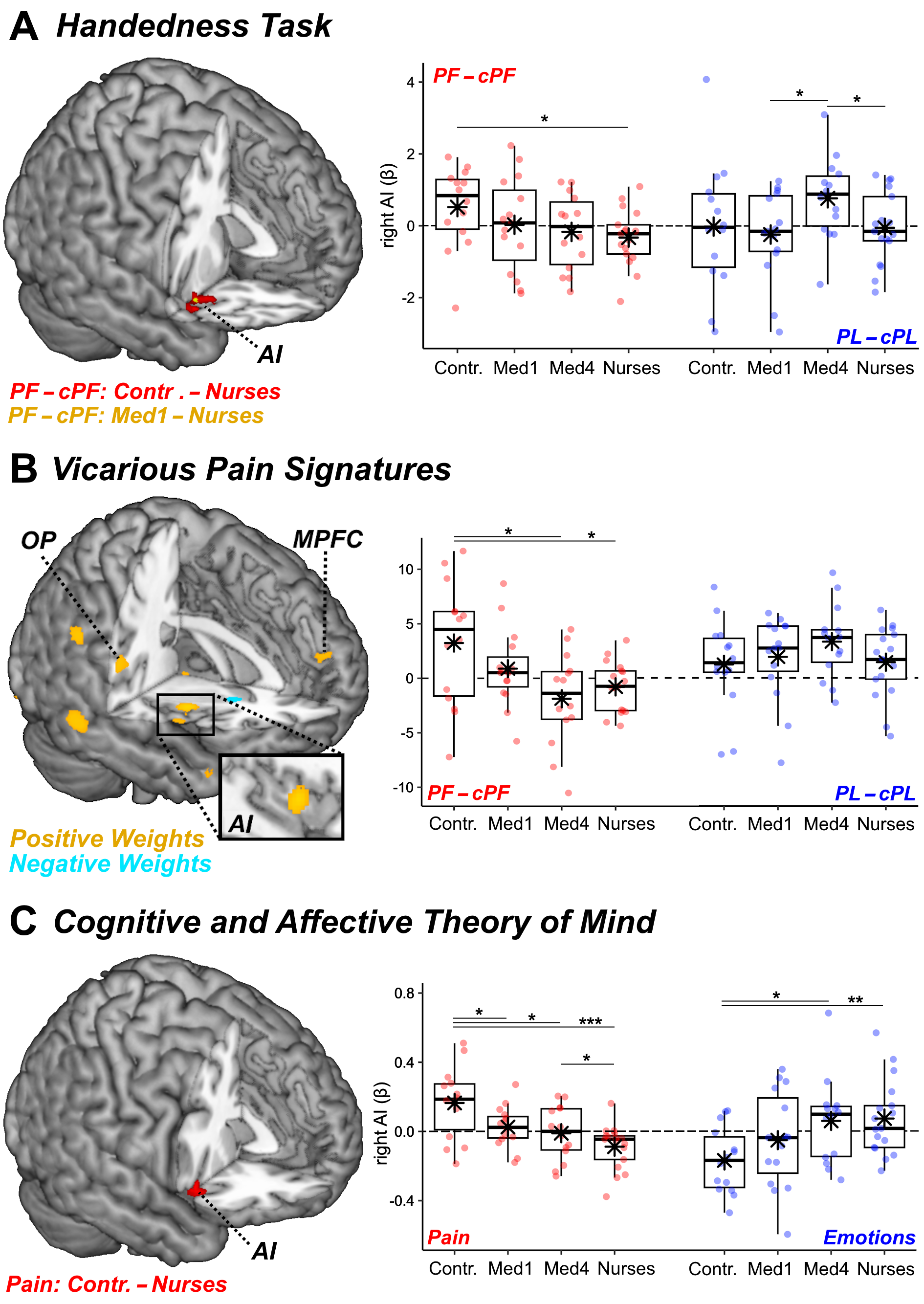
